# Mechanisms of disrupted neurodevelopment after Zika virus infection in infancy

**DOI:** 10.64898/2026.01.26.701807

**Authors:** Venkata-Viswanadh Edara, Sienna Freeman, Maureen Sampson, Kathryn Moore, Rebecca Richardson, Nils Schoof, Divine Burgess, Michelle J. Lee, Gregory Tharp, Roza Vlasova, Shane Taylor, Ariana Hodjatzadeh, Yihana Melendez-Alejandro, Samia Abdallah, Esther Ndungu, Ava Mascarenhas, Chao-Hsiung Hsu, Tsang-Wei Tu, Martin Styner, Mehul S. Suthar, Mar M. Sanchez, Steven Bosinger, Mark Burke, Steven A. Sloan, Jessica Raper, Ann Chahroudi

## Abstract

Congenital and early-life Zika virus (ZIKV) infection can result in neurologic deficits. Precise mechanisms of injury, especially in the more subtle presentation of postnatal infection, are not fully elucidated. Here, we defined the effects of ZIKV on the developing brain using single cell transcriptomics, histopathology and design-based stereology, diffusion MRI, and neurobehavioral assessments in infant rhesus macaques. ZIKV upregulated interferon-stimulated genes in activated microglia and cell death pathways in neurons and downregulated metabolism and differentiation genes in mature oligodendrocytes. Abnormal micro-organization of the corpus collosum and limbic white matter tracts was seen on diffusion weighted imaging. A curated gene set associated with autism spectrum disorder risk was negatively enriched in inhibitory and excitatory neurons from ZIKV-infected infants, with increased emotional reactivity already evident two weeks following infection. From single cells to organism-level behaviors, these results define the pathways and processes disrupted by early-life ZIKV infection.

## Introduction

Intense research has focused on the neuropathogenesis of congenital Zika virus (ZIKV) infection including microencephaly. Studies of microencephaly implicate loss of fetal intermediate progenitor cells, particularly in the subventricular zone of the developing brain(*1–4*). Progenitor cell depletion has been attributed to apoptosis, immune activation, and cell cycle dysregulation(*5–9*). Microcephaly is an extreme phenotype of congenital ZIKV infection, and more subtle features including intracranial calcifications, cerebral atrophy, and optic nerve abnormalities, also occur(*10–12*). Similarly, postnatal ZIKV infection in infants and children has been observed, with acute neurologic complications that include encephalitis and Guillain-Barré Syndrome(*13–15*). Delayed neurodevelopment following early life ZIKV infection has also been described(*16*) although challenges in diagnosis make a direct association difficult to confirm on a population level. Some but not all long-term follow up studies of children with postnatal ZIKV exposure adverse neurodevelopment, including hearing or vision loss and poor performance in the personal-social domain of a standardized developmental screening tool(*17–21*).

The large body of data demonstrating extensive and dynamic postnatal growth and maturation of the brain over the first years of life(*22, 23*) suggests that ZIKV infection in this period could be harmful and calls for definitive studies in carefully controlled model systems. Critical regions such as the hippocampus, which continue maturing postnatally and are essential for learning, memory, and behavior, are particularly vulnerable to disruption, a factor implicated in neurodevelopmental disorders such as autism and schizophrenia(*24, 25*). Given this vulnerability, understanding the effects of ZIKV infection beyond the prenatal period is essential. We previously described long-term consequences of early life ZIKV infection on the brain and behavior of infant rhesus macaques with partial mitigation by antiviral treatment(*26–28*). Rhesus macaques exhibit neurodevelopment that is temporally and anatomically comparable to humans, albeit on an accelerated timeline of approximately 1:4 ratio(*29*). In fact, transcriptomic signatures of neural processes in humans and macaques converge in the late fetal to infant (perinatal) development periods(*30*). The prolonged postnatal brain maturation, immune system similarity, and comparable social behaviors of rhesus macaques make them particularly well-suited for modeling human neurodevelopmental outcomes(*16, 31–34*).

Here, we aimed to elucidate the mechanisms of disrupted neurodevelopment by subjecting the hippocampus, amygdala, and striatum to single-cell transcriptomic analysis two weeks after ZIKV infection to dissect the cell-type-specific transcriptional changes that underlie ZIKV-associated neurodevelopmental disruption. These regions of the Central Nervous System (CNS) were selected based on our prior research that used in vivo Magnetic Resonance Imaging (MRI), resting state functional MRI, and postmortem histological examination to demonstrate ZIKV-induced abnormalities(*27, 28*). Acute postnatal ZIKV infection of infants led to CNS innate immune activation with an increased apoptotic gene signature and negative enrichment of autism spectrum disorder (ASD)-associated genes in neurons, as well as downregulation of maturation genes in oligodendrocytes. Complimentary histologic and stereologic examination confirmed these transcriptomic findings. In an additional 24 macaque infant cohort (12 controls and 12 ZIKV-infected), diffusion tensor imaging MRI after two months of ZIKV infection highlighted abnormal white matter organization in the limbic system and corpus callosum. Taken together, these findings provide plausible mechanisms for the abnormal brain structures, functions, and social behaviors that can result from early life ZIKV infection.

## Results

### ZIKV alters infant macaque CNS gene expression

To investigate the impact of early life ZIKV infection on brain development, we performed single-cell transcriptomic profiling of central nervous system (CNS) tissues from infant rhesus macaques (**Figure 1A**). ZIKV viremia peaked at 2-3 days post infection and was cleared from peripheral circulation within two weeks (**Figure 1B**). Prior work from our group has demonstrated that entry of viral RNA into the CNS occurs after peak viremia, with detection by PCR and RNAscope at 14-15 days of infection(*28*). Based on this finding, three infant RMs were infected with ZIKV at one month of life and euthanized 14 days post infection alongside three age-matched uninfected controls. Tissues from the amygdala, hippocampus, and striatum were harvested within 10 minutes of euthanasia and immediately processed to minimize post-mortem interval confounders, followed by single-cell RNA sequencing using the 10x Genomics platform. One ZIKV-infected infant RM died prematurely at 11 days post-infection and was excluded from the scRNA-Seq analysis; tissues were preserved for histological examination.

**Figure 1.**
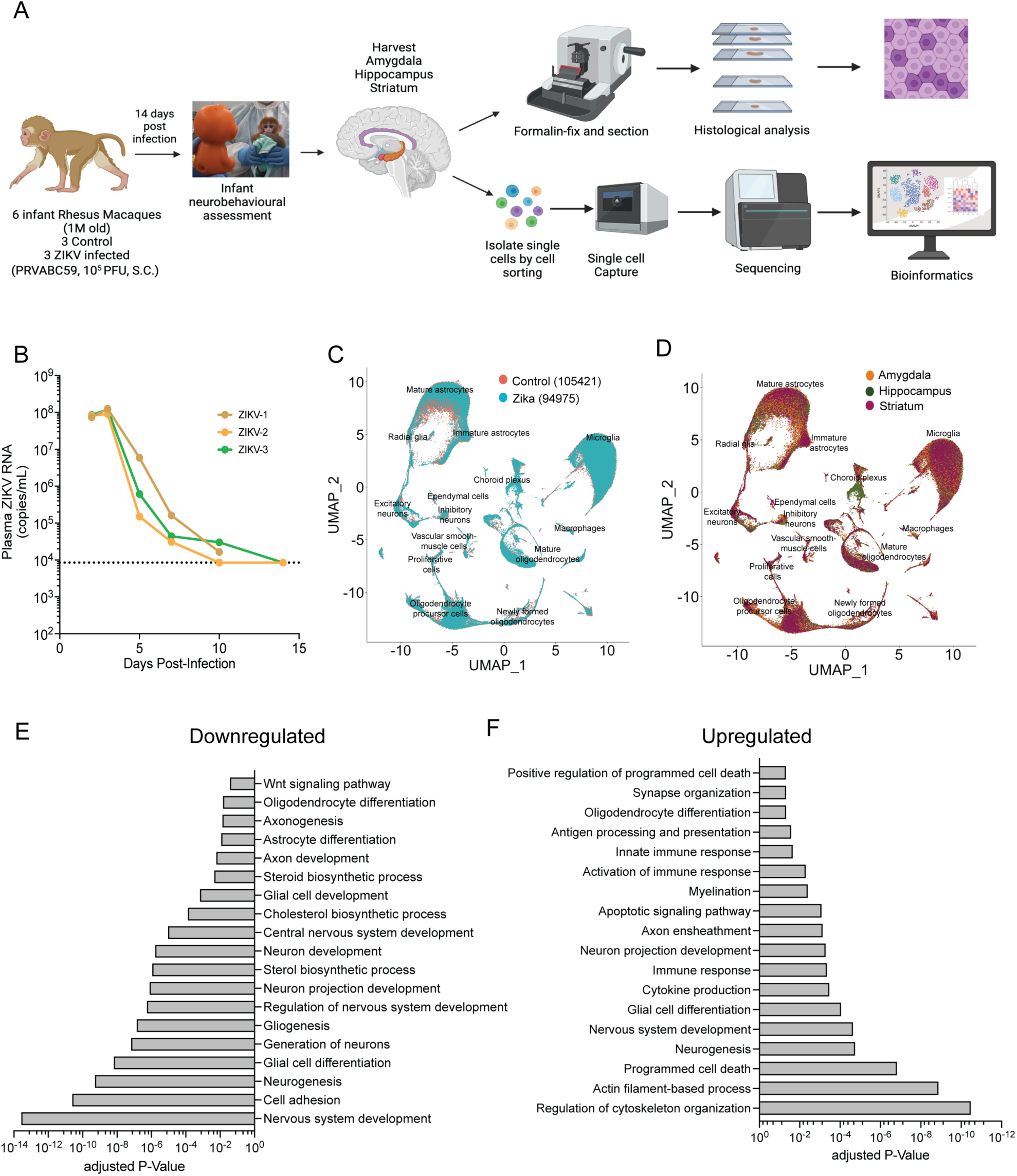
Study design, viral kinetics and central nervous system (CNS) specific transcriptome changes following ZIKV infection in infant rhesus macaques. **(A)** Study overview. Three one month old infant rhesus macaques were infected subcutaneously (s.c.) with 10^5^ plaque forming units (PFU) ZIKV strain PRVABC59. Sex and age matched uninfected animals were used as controls. Fourteen days post infection macaques were euthanized and the amygdala, hippocampus, and striatum of one hemisphere were used for single cell transcriptomic analyses and the same regions from the other hemisphere were used for histological examination. **(B)** Plasma viral loads were measured by RT-PCR from time of infection until necropsy. **(C-D)** Dimensionality reduction/UMAP analysis of combined Seurat object after quality control depicted by treatment group and brain region, respectively. **(E-F)** Gene Ontology (GO) analysis of differentially expressed genes among ZIKV-infected versus control animals showing downregulated and upregulated genes, respectively.

After quality control, 105,421 cells from controls and 94,975 cells from ZIKV-infected animals were retained for downstream analysis. Uniform manifold approximation and projection (UMAP) clustering revealed all age-relevant CNS cell classes along with their subtypes, which we subsequently re-clustered for detailed, cell-type-specific analyses. Cell type identities were assigned based on cluster-specific marker genes, with representative markers shown in the dot plot **(Supplemental Figure 1A**). UMAP plots grouped by treatment (ZIKV vs. control) and by brain region (amygdala, hippocampus, striatum) are presented in **Figure 1C** and **1D**, respectively. The distribution of cell type proportions by treatment group, brain region, and individual animal is summarized in **Supplemental Figure 1B-D**, with corresponding cell numbers shown in **Supplementary Tables 1-3**. Of all major cell types, the proportion of mature astrocytes was consistently lower in ZIKV-infected infants compared to controls, while oligodendrocyte precursor cells, mature oligodendrocytes, excitatory neurons, and ependymal cell numbers were consistently higher in ZIKV-infected animals (**Supplemental Figure 1E** and **Table 3**). The large increase in choroid plexus cells in the hippocampus was only seen in one ZIKV-infected animal, thus was not further explored.

We first assessed global transcriptional changes associated with ZIKV infection across all cell types. Therefore, we performed differential expression analysis across all cells comparing ZIKV-infected to control animals, identifying significantly differentially expressed genes (DEGs) using a false discovery rate (FDR) threshold of <0.05. Gene ontology (GO) enrichment analysis of downregulated genes revealed significant suppression of general pathways related to CNS development, including neuron and glial cell development and differentiation, axon development/ axonogenesis, cell adhesion, and sterol biosynthesis (**Figure 1E**). These findings suggest impaired maturation and regulation of key neurodevelopmental programs following ZIKV infection. Conversely, upregulated pathways in ZIKV-infected animals were predominantly associated with immune responses and apoptosis, including cytokine production, activation of immune response, innate immune responses, regulation of cytoskeleton organization, apoptotic signaling pathway and programmed cell death (**Figure 1F**). This robust neuroinflammatory response was also accompanied by upregulation of some nervous system development pathways.

Together, these data provide evidence that the immune response to postnatal ZIKV infection in the CNS is accompanied by both cell death and dysregulated neurodevelopmental trajectories.

### Activation of microglia and interferon signaling

Microglia have a complex, dual role in neurotropic viral infections. They serve as protective agents by defending against the virus but can also contribute to disease progression by promoting neuroinflammation(*35*). Given the robust upregulation of immune-related pathways observed in the global transcriptomic analysis, we next focused specifically on microglia to evaluate their response to postnatal ZIKV infection. First, we re-clustered all myeloid lineage cells and identified ten transcriptionally distinct subclusters (MG1-10) (**Figure 2A**). These subclusters were present in all three brain regions and in both control and ZIKV-infected animals (**Supplemental Figure 2A).** In total, we analyzed 57,477 microglial cells, with 27,212 cells originating from control animals and 30,265 from ZIKV-infected animals. The distribution of cell type proportions by treatment group, brain region, and individual animal is summarized in **Supplemental Figure 2B-D**, with corresponding cell numbers in **Supplementary Tables 4-6**. Only one (MG7) subcluster exhibited consistent cell number differences between ZIKV-infected infants and controls (**Supplementary Table 6**).

**Figure 2.**
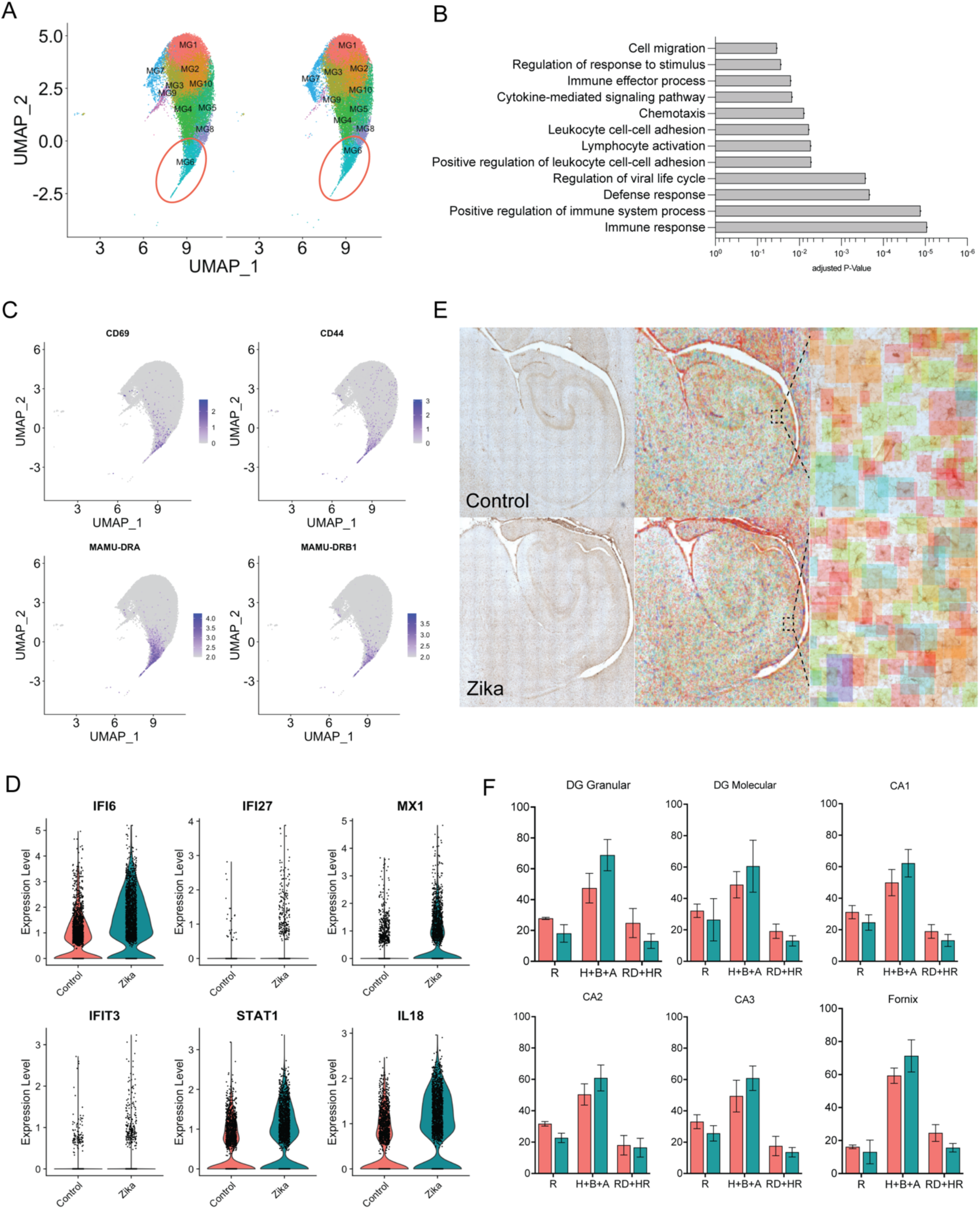
ZIKV infection leads to microglial immune activation and further induction of interferon signaling in infant rhesus macaques. **(A)** UMAP analysis of microglial subcluster segregated by treatment group. **(B)** Upregulated biological pathways from GO analysis of differentially expressed genes between ZIKV-infected versus control among microglial subcluster MG6. **(C)** Feature plots of activated microglial markers and MHC class II molecules. **(D)** Expression of interferon stimulated genes among ZIKV-infected and control microglial subcluster MG6. **(E)** IBA-1 staining of microglia from the hippocampus. Regional morphometric characterization of microglial cells was performed by Stain AI. Microglia were color-coded as ramified (green), hypertrophic (gold), bushy (orange), amoeboid (red), rod (blue), hyper-rod (purple) or unfocussed (gray). **(F)** Quantification of surveillant (ramified) versus activated (hypertrophic, bushy, and amoeboid) microglia in the hippocampal gray matter of ZIKV-infected versus control infant rhesus macaques. Mean and SD are shown. Teal bars = ZIKV; salmon bars = controls.

To identify specific microglial populations associated with immune activation, we examined gene expression patterns across all subclusters. This analysis revealed that subcluster MG6 had elevated expression of immune response genes. Using GO pathway analysis to compare groups, MG6 cells from ZIKV-infected macaques exhibited enrichment for pathways associated with immunity, including positive regulation of immune system process, defense response, lymphocyte activation, chemotaxis, cytokine-mediated signaling pathway, and immune effector process (**Figure 2B**). Cluster MG6 cells in ZIKV-infected infants showed elevated expression of canonical myeloid cell activation markers CD69, CD44, and MHC class II molecules MAMU-DRA and MAMU-DRB1 (**Figure 2C**). To comprehensively assess interferon pathway activation, we interrogated the differential expression of a panel of interferon-stimulated genes within both the entire microglial population and the MG6 subcluster in the combined dataset and across individual brain regions (**Supplemental Figure 3A** and **3B**, respectively). These analyses revealed widespread and robust microglial induction of interferon stimulated genes in response to ZIKV infection. Elevated expression of representative interferon stimulated genes IFI6, IFI27, MX1, IFIT3, STAT1, and IL18, specifically within the MG6 subcluster, is shown in **Figure 2D**.

**Figure 3.**
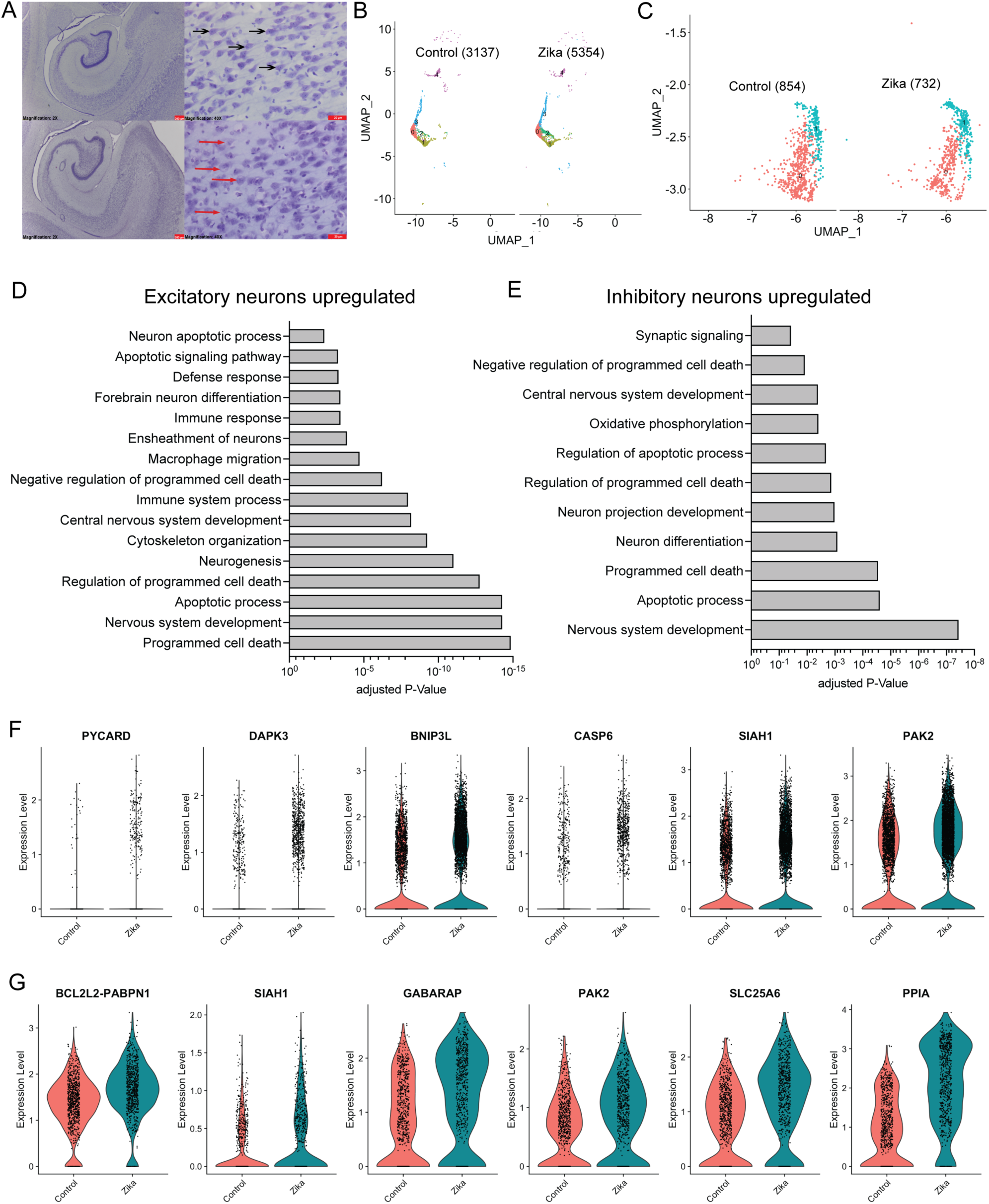
ZIKV infection leads to induction of apoptotic pathways in the limbic region of infant rhesus macaques. **(A)** Cresyl violet stained mature neurons in the hippocampal CA1 subfield showing frequent cytoplasmic inclusions (red arrows) in ZIKV-infected macaques and healthy neurons (black arrows) in control macaques. Top panels are representative images from control and bottom panels are from ZIKV-infected animals (scale bar left panels=200 µm; scale bar right panels=20 µm). **(B)** UMAP analysis of excitatory neuron subcluster segregated by treatment group. **(C)** UMAP analysis of inhibitory neuron subcluster segregated by treatment group. **(D)** Upregulated biological pathways from GO analysis of differentially expressed genes between ZIKV-infected versus control animals among excitatory neurons. **(E)** Upregulated biological pathways from GO analysis of differentially expressed genes between ZIKV-infected versus control animals among inhibitory neurons. **(F)** Expression of selected cell death genes in excitatory neurons from ZIKV-infected versus control macaques. **(G)** Expression of selected cell death genes in inhibitory neurons from ZIKV-infected versus control macaques.

We next used design-based stereology and histological examination of hippocampi to identify microglia by Iba1 staining. Microglia were subcategorized into ramified, hypertrophic, bushy, amoeboid, rod, and hyper-rod based on their morphologic features in the following regions of the hippocampal formation: dentate gyrus granular and molecular layers, *Cornu Ammonis* (CA) subfields 1-3, and fornix. Iba-l staining was quantified for each cell type using machine learning techniques (**Figure 2E**). The frequency of Iba-1+ activated microglia in the grey matter (hypertrophic, bushy, and amoeboid cells) was increased in ZIKV-infected infants compared to controls across all six regions of the hippocampus (**Figure 2F**). Although the small number of animals precluded formal statistical analysis, this difference was most pronounced in the granular region of the dentate gyrus, where microglia regulate neurogenesis through phagocytosis of dying granule cells(*36, 37*) as part of a homeostatic process involved in memory formation, pattern separation, and spatial behavior(*38–40*). These collective findings suggest that postnatal ZIKV infection induces a shift in the microglial population toward an activated, interferon-responsive state, potentially influencing neurogenesis.

### ZIKV upregulates neuronal apoptosis pathways

Design-based stereology was used to quantify hippocampal neuronal populations of the pyramidal layer in CA subfields 1-3 in cresyl violet stained sections and nestin positive immature neurons in the dentate gyrus. Counts were mostly similar between ZIKV-infected and control infants, with a trend towards reduced CA1 neurons. (**Supplemental Figure 4A-B**). Notably, mature neurons in the vulnerable CA1 region of ZIKV-infected macaques showed frequent cytoplasmic inclusions, typically a precursor to cell death, that was not seen in controls (**Figure 3A**). This finding is consistent with the upregulation of apoptosis pathways seen in the initial global transcriptomic analysis (**Figure 1F**). To evaluate how postnatal ZIKV infection influences neuronal subtypes critical for cortical circuit development, we next examined gene expression in both excitatory and inhibitory neuronal populations (a total of 10,077 neurons were studied: 3,137 excitatory and 854 inhibitory neurons from controls and 5,354 excitatory and 732 inhibitory neurons from ZIKV-infected macaques; (**Figure 3B-C**). Region-specific distributions of these neuronal populations across the amygdala, hippocampus, and striatum as well as cell proportions by treatment, region, and individual animal, are shown in **Supplemental Figure 5**, with corresponding cell numbers in **Supplementary Tables 7-12**. GO pathway analysis revealed upregulation of multiple cell death pathways in both neuron subtypes in ZIKV-infected macaques (**Figure 3D-E**). Both excitatory and inhibitory neurons of ZIKV-infected infants compared to controls showed increased expression of genes involved in regulating apoptosis (PYCARD, DAPK3, BNIP3L, CASP6, SIAH1, PAK2, BCL2L2-PABPN1, PPIA), autophagy (GABARAP), and mitophagy (SLC25A6) (**Figure 3F-G**). These results indicate that induction of cell death signaling pathways is a common feature of postnatal and congenital ZIKV infection, with anticipated consequences for neurodevelopment.

### Neurodevelopmental transcriptional program in neurons and behavior changes

We next assessed whether ZIKV-induced gene expression changes in neuronal populations resembled shifts observed in other neurodevelopmental disorders. To do so, we interrogated the transcriptomic signature of neurons using a curated panel of 147 genes(*41–45*) implicated in the common neurodevelopmental disorder, autism spectrum disorder (ASD), and categorized them based on their expression timing in Epoch-1 (prenatal) and Epoch-2 (late prenatal and early postnatal) phases of neurodevelopment(*41, 46*). In excitatory neurons, GSEA revealed downregulation of both Epoch-1 and Epoch-2 gene sets in ZIKV-infected macaques (enrichment scores = -0.372 and -0.500, respectively), with a significant net enrichment score for Epoch-2 (NES = -1.49, *p* = 0.0451; **Figure 4A**). In inhibitory neurons, both Epoch-1 and Epoch-2 gene sets were similarly downregulated (enrichment scores = -0.461 and -0.429), with the Epoch-1 gene set demonstrating statistical significance (NES = -1.50, *p* = 0.0089; **Figure 4B**). To test whether these gene expression changes were nonspecific, we performed GSEA using two independent gene sets associated with cerebral palsy. These analyses did not reveal significant associations with ZIKV infection (**Supplemental Figure 6A-D**). Further, we tested the enrichment of these ASD associated gene sets in other neural cell populations, including mature oligodendrocytes (**Supplemental Figure 6E-F**), oligodendrocyte progenitors, and microglia, none of which showed significant enrichment. These control analyses support the specificity of ASD gene dysregulation in excitatory and inhibitory neurons in the context of ZIKV infection. High-confidence ASD risk genes, including ANK2, CHD8, CUL3, DYRK1A, GRIN2B, KATNAL2, POGZ, SCN2A, and TBR1, were consistently downregulated in ZIKV-infected infants in both excitatory (**Figure 4C**) and inhibitory neurons (**Figure 4D**), suggesting that postnatal ZIKV infection broadly affects core neurodevelopmental pathways that control social interaction and communication.

**Figure 4.**
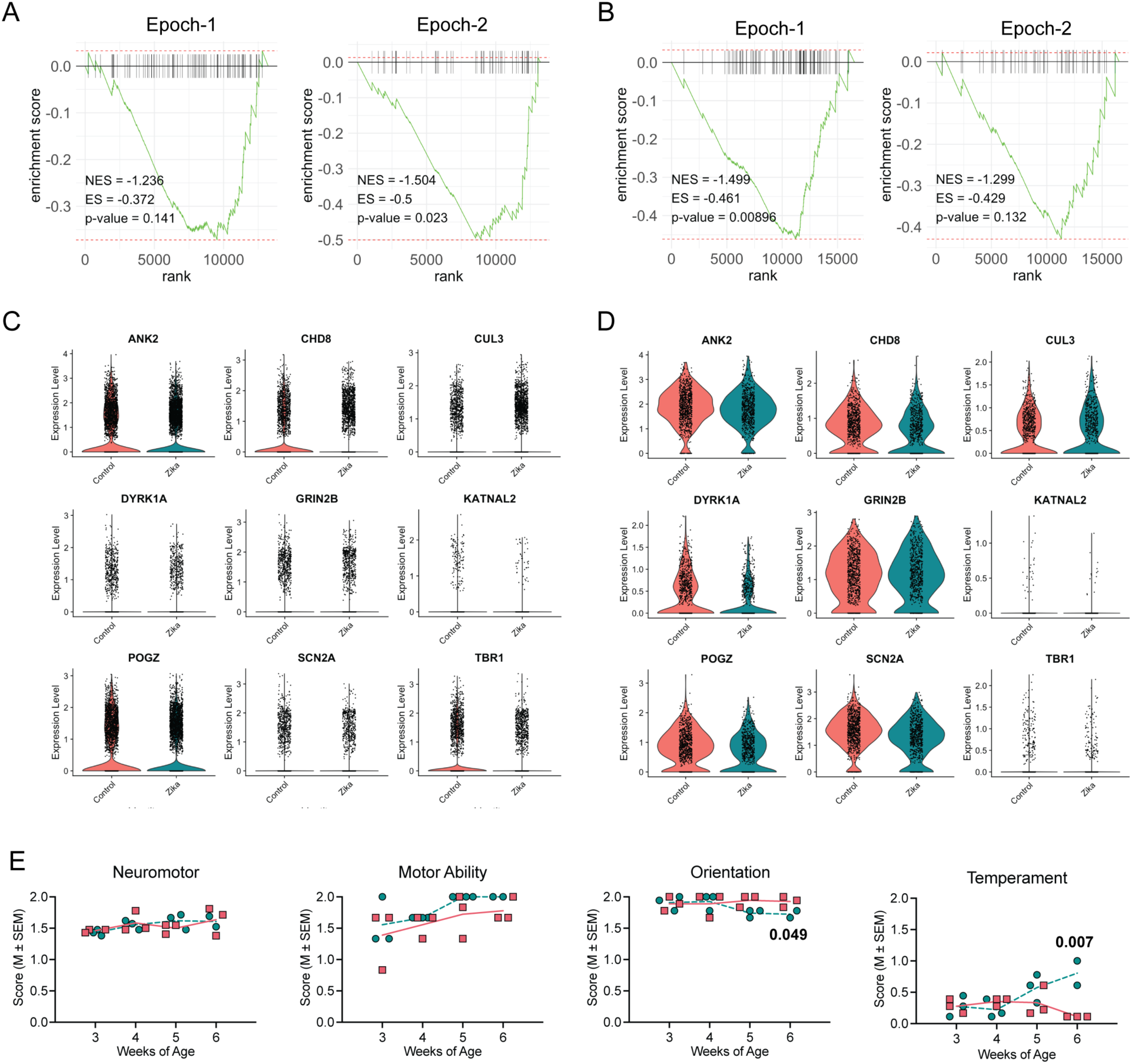
ZIKV infection leads to dysregulation of Autism Spectrum Disorder (ASD) genes among neuronal populations and neurobehavioral changes in infant rhesus macaques. **(A)** GSEA plots for a panel of 147 ASD genes spanning the Epoch-1 (prenatal) and Epoch-2 (late prenatal and early postnatal) phases of neurodevelopment in excitatory neurons of ZIKV-infected infant macaques. NES, normalized enrichment score. ES, enrichment score. **(B)** GSEA plots for 147 ASD genes spanning the Epoch-1 (prenatal) and Epoch-2 (late prenatal and early postnatal) phases of neurodevelopment in inhibitory neurons of ZIKV-infected infant macaques. **(C)** Expression of high confidence ASD genes in excitatory neurons of ZIKV-infected and control infants. **(D)** Expression of high confidence ASD genes in inhibitory neurons of ZIKV-infected and control infants. **(E)** Battery of neurobehavioral tests assessing neuromotor skills, motor ability, orientation and temperament in ZIKV-infected versus control infant macaques from 3 to 6 weeks of life. Differences were assessed using a linear mixed model with group and age as fixed factors and individual animals as a random factor. Teal symbols/lines = ZIKV; salmon symbols/lines = controls.

To relate these molecular findings to functional outcomes, we examined early neurobehavioral development in the same infant animals. We have previously shown that postnatal ZIKV infection at one month of life results in reduced prosocial behaviors and increased emotional reactivity at one year(*27*). Here, at a very early age, postnatal ZIKV infection was associated with a significant decrease in orientation scores (*F*[3,21] = 3.60, *p* = 0.049, ηp² = 0.34) and an even more striking increase in temperament scores (*F*[3,21] = 6.84, *p* = 0.007, ηp² = 0.49) (**Figure 4E**). Both groups had similar neuromotor scores and typical age-related increases in motor ability from 3 to 6 weeks. Early alterations in attention and behavioral regulation, as seen here, are phenotypes frequently reported in children with neurodevelopmental disorders(*47–50*).

### Disruption of differentiation and metabolic programs in oligodendrocytes

Among all the differences observed between ZIKV-infected and control infant macaques, oligodendrocyte differentiation was one of the most prominent changes. Oligodendrocytes are glial cells in the central nervous system responsible for forming myelin sheaths around axons, which insulate the fibers and enhance the speed of electrical signal transmission(*51*). During brain development, oligodendrocyte precursor cells differentiate into mature oligodendrocytes to create these sheaths, a process essential for building functional neural networks and facilitating sensory information processing(*51*). To assess how postnatal ZIKV infection affects oligodendrocyte lineage cells, we performed focused sub-clustering of oligodendrocytes from single-cell transcriptomic data. From a total of 51,899 cells (22,168 from controls and 29,731 from ZIKV-infected macaques), UMAP analysis revealed three major populations: oligodendrocyte progenitor cells (OPCs), newly formed oligodendrocytes, and mature oligodendrocytes (**Figure 5A**). Subpopulation identity was determined through examination of canonical lineage marker expression, with enrichment of OLIG2 in OPCs, BCAS1 in newly formed oligodendrocytes, and MBP in mature oligodendrocytes (**Figure 5B**). The distribution of these subclusters in the amygdala, hippocampus, and striatum, with proportions by treatment group, brain region, and individual animal are depicted in **Supplemental Figure 7**. Corresponding cell numbers are shown in **Supplementary Tables 13-15**. The ZIKV-infected macaques had increased proportions of both OPCs and mature oligodendrocytes compared to controls (**Supplementary Table 13**).

**Figure 5.**
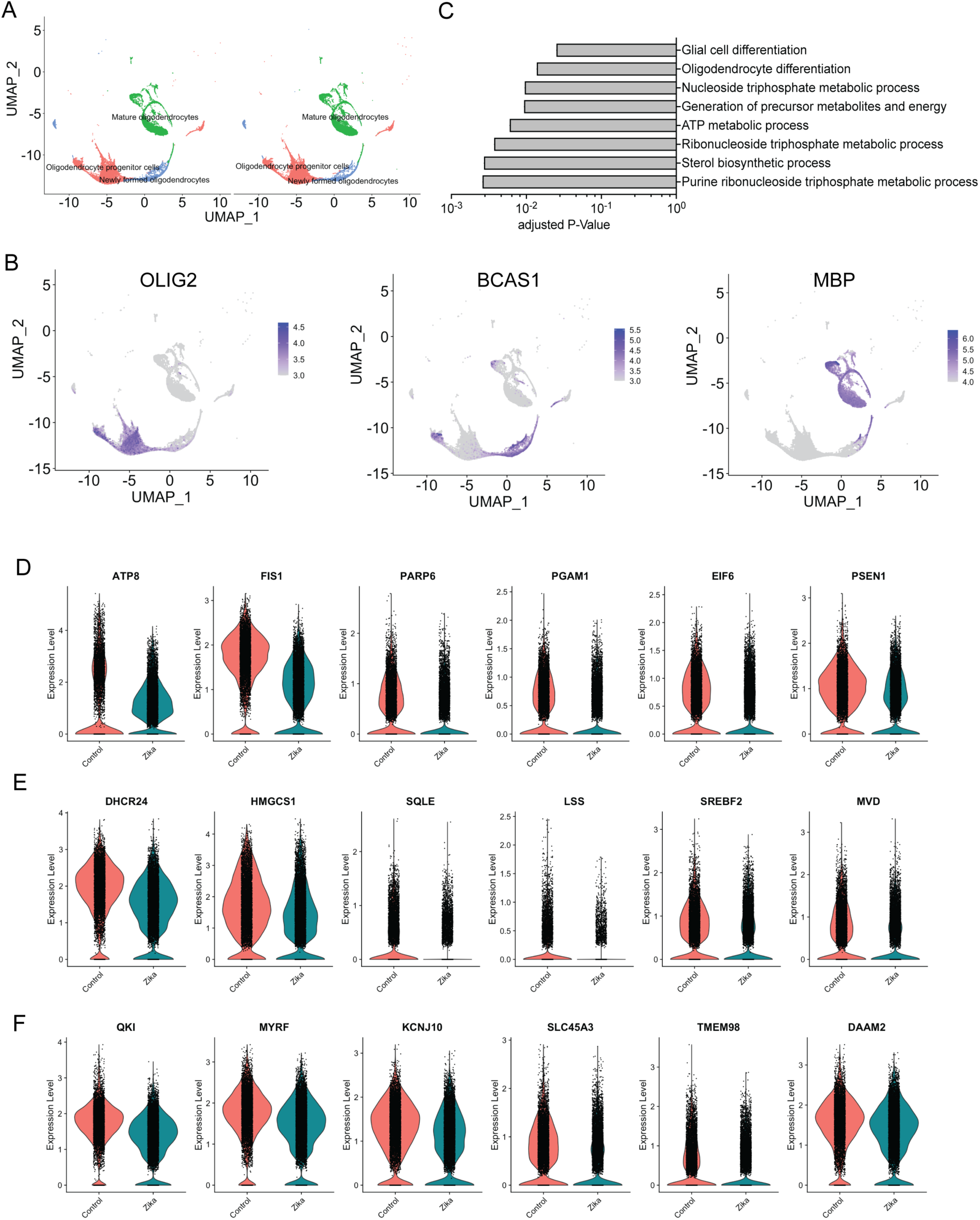
ZIKV infection dysregulates key biological pathways in mature oligodendrocytes of infant rhesus macaques. **(A)** UMAP analysis of oligodendrocyte subclusters segregated by treatment group. **(B)** Feature plots of lineage markers OLIG2, BCAS1, and MBP used to distinguish oligodendrocyte progenitor cells, newly formed oligodendrocytes, and mature oligodendrocytes, respectively. **(C)** Downregulated biological pathways from GO analysis of differentially expressed genes between ZIKV-infected versus control mature oligodendrocytes. **(D)** Expression of representative genes regulating metabolic processes in mature oligodendrocytes from ZIKV-infected and control infant macaques. **(E)** Expression of selected genes involved in sterol biosynthesis in mature oligodendrocytes from ZIKV-infected and control infant macaques. **(F)** Expression of genes controlling oligodendrocyte differentiation in mature oligodendrocytes from ZIKV-infected and control infant macaques.

To determine how ZIKV infection alters mature oligodendrocyte function, we performed differential gene expression and GO analysis between ZIKV-infected and control oligodendrocyte populations. Downregulated pathways (lower in ZIKV) were significantly enriched for terms related to metabolic processes, sterol biosynthesis, and oligodendrocyte and glial cell differentiation (**Figure 5C**). Volcano plots of representative genes from these pathways further highlight their suppression in ZIKV-infected animals (**Supplemental Figure 8**).

Expression of individual genes involved in cellular metabolism (ATP8, FIS1, PARP6, PGAM1, EIF6, PSEN1) (**Figure 5D**), sterol biosynthesis (DHCR24, HMGCS1, SQLE, LSS, SREBF2, MVD) (**Figure 5E**), and oligodendrocyte differentiation and maturation (QKI, MYRF, KCNJ10, SLC45A3, TMEM98, DAAM2) (**Figure 5F**) were all significantly reduced following ZIKV infection. These data indicate that postnatal ZIKV infection perturbs oligodendrocyte development by disrupting core metabolic pathways, suppressing sterol biosynthesis, and impairing terminal differentiation—processes that are critical for CNS maturation and myelination during early development(*52–56*). Concurrent histologic staining of the corpus callosum and limbic system white matter tracts (cingulum, fornix) showed similar myelin levels in infants with or without ZIKV infection (**Supplemental Figure 9**) indicating that the transcriptomic changes we observed in oligodendrocytes after two weeks of ZIKV infection did not simultaneously impact myelin density in these regions.

### Abnormal white matter micro-organization at 3 months of age

To evaluate white matter microstructure and longer-term impact of the CNS transcriptomic changes induced by postnatal ZIKV infection, we followed an additional 24 macaque infants (12 controls and 12 ZIKV-infected) until three months of age. These infants underwent MRI with diffusion tensor imaging (DTI) of the brain to investigate white matter micro-organization.

Fifteen white matter fiber tracts were reconstructed in a high-resolution study-specific DTI atlas and fractional anisotropy (FA), axial diffusivity (AD), mean diffusivity (MD), and radial diffusivity (RD) were sampled along each tract’s arc length. **Figure 6A** shows representative examples of the corpus callosum dorsal segment 1 (cc1d), cingulum (cg), and fornix in a gray matter surface of a macaque brain in DTI atlas space. We next compared the mean FA, AD, MD, and RD for the ZIKV-infected versus control infants, focusing on the largest white matter tract (corpus callosum) and tracts projecting between limbic system structures. FA was increased in the ccld, right cingulum (cg_r; projecting from the cingulate gyrus to entorhinal cortex), and right fornix (major output tract of the hippocampus) of ZIKV-infected macaques (**Figure 6B-D**), indicating abnormal white matter micro-organization. This increase in FA was paralleled by increased AD with similar mean MD and RD compared to controls. This pattern can be a sign of accelerated white matter maturation, consistent with inflammation-induced axonal damage accompanied by repair-triggered enhanced myelination(*57, 58*). Comparisons of FA for all 15 white matter tracts between ZIKV-infected versus control infants as well as within each group by sex are shown in **Supplementary Table 16**. In conclusion, these data reveal mechanisms of disrupted neurodevelopment that involve CNS immune activation, programmed cell death, cortical circuit regulation, and oligodendrocyte maturation resulting in microstructural changes to white matter and increased emotional reactivity.

**Figure 6.**
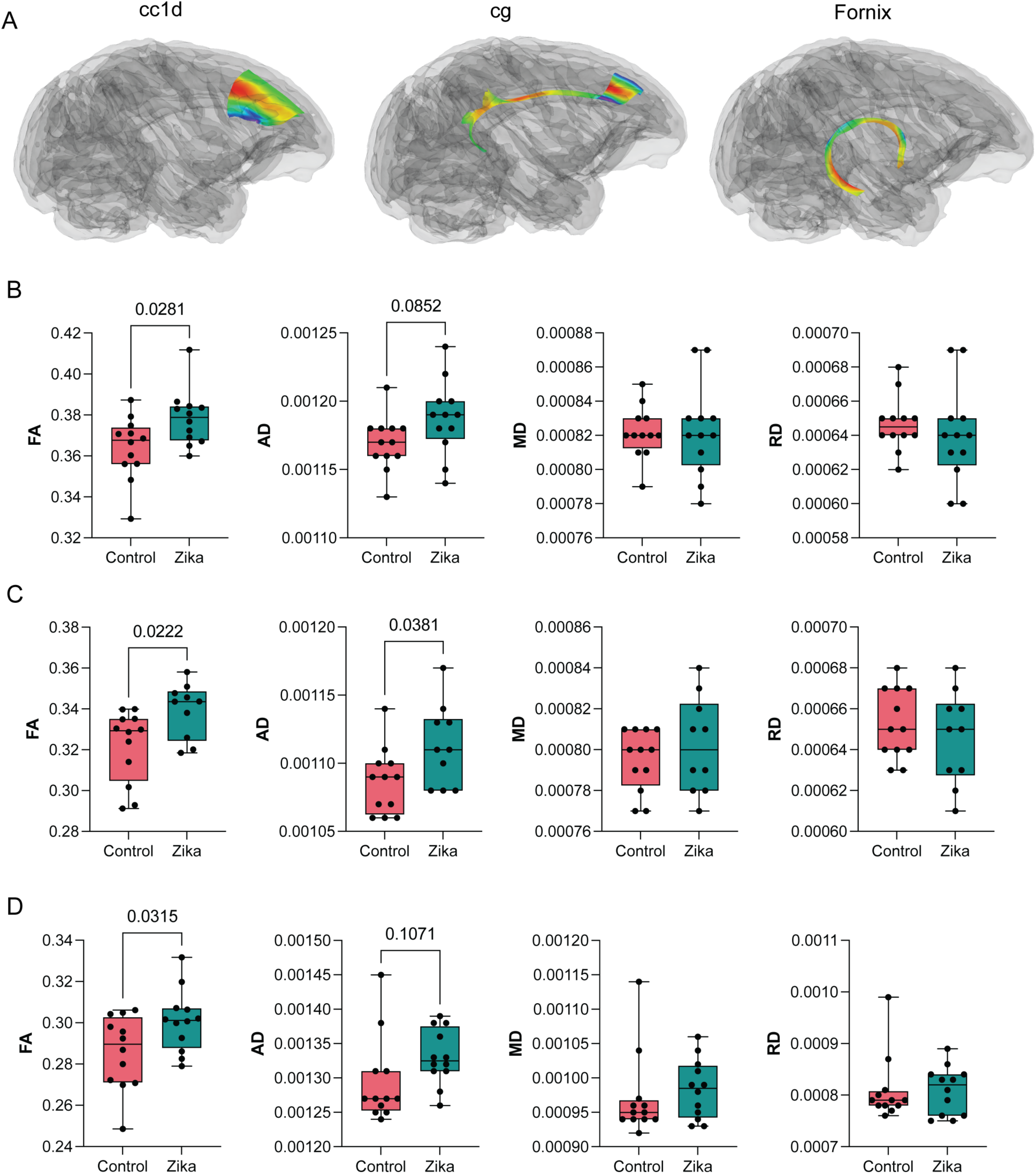
Postnatal ZIKV infection alters corpus callosum and cingulate gyrus micro-organization. **(A)** Diffusion tensor images (DTI) of white matter tracts of corpus callosum dorsal segment 1 (cc1d), cingulum (cg) and fornix visualized by projecting onto a gray matter surface in DTI atlas space. **(B)** Fractional anisotropy (FA), axial diffusivity (AD), mean diffusivity (MD), and radial diffusivity (RD) of ZIKV-infected versus control infants for cc1d. **(C)** FA, AD, MD, and RD of ZIKV-infected versus control infants for the right cg. **(D)** FA, AD, MD, and RD of ZIKV-infected versus control infants for the right fornix. Two-factor ANOVA was used to investigate effects of group (ZIKV and control) and interaction term on FA, AD, RD, and MD. Boxes represent the first to third percentiles, with the central line showing the median and the whiskers indicating the minimum and maximum values.

## Discussion

The specific set of ZIKV-induced changes in the CNS transcriptome we report is highly suggestive of impaired neurodevelopment following early life ZIKV infection. Results of neuroimaging and behavioral testing performed in this study and prior research show longer-term brain structural and functional abnormalities resulting from postnatal exposure to ZIKV(*27, 28*). Human infants and children infected with ZIKV do not display the more severe anatomic deformities that encompass congenital ZIKV syndrome (CZS)(*59, 60*), though developmental disabilities have been noted(*17*), similar to reports of normocephalic newborns with congenital exposure(*61, 62*). Here, using ZIKV infection of four-week-old rhesus macaque infants, we demonstrate massive divergence from uninfected control macaques, with aberrant microglial activation, neuronal apoptosis and reduced expression of genes governing neurodevelopment, as well as altered oligodendrocyte maturation and disorganized myelin tracts.

The ZIKV outbreak of 2015-2016 in Central and South America is believed to have resulted in over 700,000 infections(*63*). With ZIKV now endemic in many countries, ongoing infection is assured, although detection outside of pregnancy is more variable. With this backdrop and considering the low to middle income status of countries with ZIKV in circulation, an epidemiologic assessment of the onset of developmental disability following early life ZIKV infection is next to impossible. Prospective cohort and observational studies have shown delayed neurodevelopment in some countries but not others(*17–21*). Additionally, the multiple descriptions of congenital ZIKV infection resulting in postnatal onset of microcephaly(*64–66*) further reinforce the dynamic and critical nature of early life brain development. Therefore, the NHP model employed here provides an important system to better understand the consequences of ZIKV infection in infancy.

We have previously shown virus dissemination into the central and peripheral nervous system by two weeks after infection using PCR and RNAscope(*28*). It is likely that direct infection plays a role in the negative outcomes of ZIKV infection, but the consequences of the antiviral immune response and immune activation may also be substantial. To that end, we observed that postnatal ZIKV infection induced a subset of activated microglia characterized by upregulation of transcription factors involved in interferon signaling, leading to increased expression of interferon stimulated genes including *IFI6*, *IFI27*, and *MX1*. A similar immune activation signature is seen in other neurotropic viruses, such as West Nile, Dengue, Herpes Simplex, and HIV(*67, 68*), where neuroinflammation causes significant pathology. Interestingly, a role for type I interferon, specifically IFN-β, in regulating ZIKV-induced apoptosis in human fetal brain explants has recently been proposed(*69*). In congenital ZIKV infection, virus triggered apoptosis of fetal intermediate progenitor cells has been associated with ventricular enlargement and microcephaly(*6–9, 70*). We did not identify these neural progenitor cells in our single cell preparations, likely due to the macaque age at the time of necropsy. However, we uncovered an apoptosis signature in both excitatory and inhibitory neurons, with histologic signs of apoptosis in hippocampal neurons. Collectively, these findings indicate that loss of neurons in the developing brain is a hallmark of both congenital ZIKV syndrome and postnatal ZIKV infection and that innate immune activation may both limit infection and promote immunopathology.

In terms of linking gene expression data to potential outcomes, we noted downregulation of genes expressed by neurons that are known to be associated with ASD risk(*41*). The specificity of this result is highlighted by the fact that CP gene sets were not differentially expressed in ZIKV-infected macaques and that only neurons exhibited the ASD transcriptional program.

Together with the observation of increased emotional reactivity on behavioral testing and our prior finding of abnormal social interactions(*27*) these data support a link between altered socioemotional behavior and the specific genes linked to increased ASD risk. Thus, we postulate that there is potential for an ASD-like neurodevelopmental disorder to arise in some individuals exposed to ZIKV in early life. Poor personal-social performance on a standardized developmental screening tool has in fact been documented in children with ZIKV exposure(*17*). Our analysis was limited by the nature of RNA-Sequencing that permits an understanding of the relative levels of gene expression but not specific variants or epigenetic factors, and it should be recognized that many genes linked to ASD risk are not merely downregulated but instead have mutations or are loss of function variants(*71, 72*).

Investigation of white matter tracts by MR DTI in three-month-old macaques demonstrated the ongoing impact of prior ZIKV infection at one month of age. Somewhat surprisingly, given the downregulation of oligodendrocyte metabolism and differentiation genes at two weeks post infection, at 3 months ZIKV-infected infants had evidence of accelerated white matter maturation in the cingulum, fornix, and corpus callosum. Increased FA, as we found with ZIKV, can be seen in some white matter tracts in following preterm birth(*73–75*) and in adolescents(*76*) with severe perinatal brain injury as well as in the setting of mild cognitive impairment(*77, 78*) and mild traumatic brain injury(*79*). One interpretation of these findings is that a repair-driven compensatory response to the viral insult results in disorganized myelination of white matter.

Intriguingly, in a pigtail macaque model of congenital infection, ZIKV was associated with downregulation of genes regulating myelination and myelin decompaction in the deep white matter(*80*). Thus, both age at infection and duration of follow-up after infection are important factors when interpreting the structural implications of differential gene expression, particularly in light of the remarkable plasticity of the brain.

Using a clinically relevant infant nonhuman primate model, this research provides novel mechanistic insight into the cells and gene signatures that are modulated by ZIKV infection with indications as to how this response disrupts the typical neurodevelopmental trajectory. We show that postnatal ZIKV infection of four-week-old rhesus macaques has a distinct, nuanced, and mutable impact on the brain, underscoring the potential vulnerability of newborns and infants with ZIKV exposure.

### Limitations of the study

There were limitations to this work. First, a limited number of animals were subjected to transcriptomic analysis. This limitation was partially mitigated by the large number of single cells studied and the analysis of three separate brain regions. ZIKV infection led to relatively consistent changes across the amygdala, hippocampus, and striatum which may be related to a global viral impact and/or technical limitations with single cell preparations that may cause data homogenization. Future studies utilizing spatial transcriptomic approaches would allow for *in situ* gene expression profiling with a deeper examination of regional cell organization and interaction. In addition, we note that studying gene expression does not capture the impact of epigenetic and post-translational modifications that influence protein activity. Multi-omic technologies coupled with longer term behavioral and cognitive assessments will likely provide a more complete understanding of postnatal ZIKV infection.

## Methods

### Animals

Thirty infant Indian RMs (*Macaca mulatta*), thirteen males and seventeen females, were included in this study. Fifteen infants were infected with ZIKV at four weeks of age and fifteen infants served as age-, sex-, and rearing-matched uninfected controls. The infants were delivered naturally by their dams while housed in indoor/outdoor social groups at the Emory National Primate Research Center (ENPRC). All dams were from the specific pathogen-free colony (negative for Herpes B, SIV, SRV, and STLV1), had not been previously used for infectious disease or vaccine studies, and did not have any clinical signs of infection during pregnancy. Infants were then removed from their dams at 12-16 days of age and transported to the ENPRC nursery facility. All infants rotated daily between being singly housed or pair housed in warming incubators for the first 3-4 weeks. When single housed, infants still had visual and auditory contact with conspecifics. Infants were hand-fed formula every 2–3 h for the first month, then via self-feeders for the remainder of the study, as per standard ENPRC protocol. Soft blankets, plush toys, or fleece were provided and changed daily. Soft chow (Purina Primate Chow, Old World Monkey formulation) and fruits were introduced starting at 1 month of age.

Water was provided ad libitum. At 5 weeks of age, infants transitioned into age-appropriate caging and housed in trios consisting of full tactile, visual, and auditory contact with conspecifics. During the ZIKV-infected period, infants were housed in an ABSL-2+ (Animal Biosafety Level 2 enhanced) room until virus was no longer detected in blood or urine. All housing was indoors on a 12 h light–dark cycle.

### Virus and infection

The ZIKV of Puerto Rican origin (PRVABC59, GenBank accession number: KU501215.1) used in this study had been passaged four times and titrated on Vero cells. Experimental infections were performed via the subcutaneous route using 10^5^ plaque forming units (pfu) of ZIKV PRVABC59. One ZIKV-infected infant RM died at 11 days post-infection and had to be excluded from scRNA-Seq analysis, although tissues were preserved for histological examination. The cause of death was determined to be lung congestion due to pneumonia. PCR of lung tissues was negative for SARS-CoV-2 and positive for ZIKV.

### ZIKV RNA detection by qRT-PCR

Total RNA was extracted from 140 μl of plasma and CSF samples using the QIAamp Viral RNA Mini Kit (Qiagen). RNA from urine was isolated using the QIAamp Circulating Nucleic Acid Kit (Qiagen) for the extraction of larger volumes (up to 2 ml). ZIKV RNA standard was generated by annealing two oligonucleotides spanning the target ZIKV prM-E gene region and performing in vitro transcription using the MEGAscript T3 Transcription Kit (Ambion). Purified RNA was reverse transcribed using the High-Capacity cDNA Reverse Transcription Kit (Applied Biosystems) and random hexamers. For quantitation of viral RNA, a standard curve was generated using 10-fold dilutions of ZIKV RNA standard, and qRT-PCR was performed using TaqMan Gene Expression Master Mix (Applied Biosystems) and ZIKV primers (1 μM) and probe (250 nM). The ZIKV primer-probe set targeting the prM-E gene region included forward primer ZIKV/PR 907 (5′-TTGGTCATGATACTGCTGATTGC-3′), reverse primer ZIKV/PR 983c (5′- CCTTCCACAAAGTCCCTATTGC-3′), and probe ZIKV/PR 932 FAM (5′-CGGCATACAGCATCAGGTGCATAGGAG-3′). The probe was labeled with 6-carboxyfluorescein (FAM) at the 5′ end and two quenchers, an internal ZEN quencher and 3′ end Iowa Black FQ (Integrated DNA Technologies). The standard curve had an R2 value greater than 0.99. Viral RNA copies were interpolated from the standard curve using the sample CT value and are represented as copies per milliliter of plasma.

### Neurobehavioral Assessment

Neurobehavior was evaluated with a well-validated assessment developed for infant macaques, the Infant Neurobehavioral Assessment Scale [INAS], adapted from Schneider and Suomi(*81–83*), which is based on the human Brazelton Newborn Behavioral Assessment Scale(*84*). Twenty-nine test items in the INAS aligned with the neurodevelopmental areas of interest and make up the Orientation, Motor maturity and activity (ability), Neuromotor (sensory) responsiveness, and Temperament constructs. This neurobehavioral test has previously been used to define neonatal development of prenatally ZIKV-exposed infants(*85, 86*). Ratings were based on a five-point Likert scale ranging from 0 to 2. The INAS was administered weekly between 3-6 weeks of age. Three examiners (S.F., J.R., R.R.) were present for all neurobehavioral testing and scoring to ensure test administration reliability (> 90%). Items were administered in a consistent sequence across all animals to optimize performance and decrease handling time. Assessments were hand-scored on a printout of the scoring form during administration. Higher scores reflect optimal scores for Orientation, Motor, and Neuromotor, whereas higher Temperament scores reflect more emotional reactivity.

### Brain dissection for transcriptomics and histology

After 14 days post-infection, the animals received an overdose of pentobarbital and were transcardially perfused with phosphate buffer solution (PB; 0.2 m, pH 7.4). After removing the brain from the skull, the right and left hemispheres were immediately dissected (within 5 mins). The left hemisphere was placed in 4% paraformaldehyde until further processing, then blocked into 2 cm slabs, cryoprotected in graded buffered sucrose solutions (10-30%) and frozen in isopentane. Blocks of tissue were then systematically sectioned (50 µm) in ten parallel series with the first series of slides for cresyl violet staining and the remaining 9 series placed into antigen preserve (1% polyvinyl pyrrolidone, 50% ethylene glycol in 0.1 M PBS, Ph 7.4)(*87, 88*). The right hemisphere was systematically dissected to remove the entire amygdala, caudate, and hippocampus for transcriptomic analysis. Additionally, tissue sections from the right hemisphere were flash frozen for additional analyses.

### Histological Analysis

#### Immunohistochemistry

Series of sections were removed from antigen preserve, washed 3 times in PBS, and pretreated in 20% methanol, 3% hydrogen peroxide (in 0.1 M PBS) for 30 min at room temperature. Sections then were washed in 0.1 M PBS, blocked in a 3% normal horse serum (NHS) in PBS for 1 hour and underwent an additional set of PBS washes. Sections were then incubated overnight in either rabbit anti-Iba-1 (1:1000; Wako) or rabbit anti-nestin (Millipore, 1:1000) in the blocking solution. The following day, sections underwent a series of PBS washes, incubated in biotinylated horse anti-rabbit IgG (1:200, Vector), washed again in PBS and then incubated in ABC (Vector). The sections were then visualized with diaminobenzidine (DAB, Sigma), mounted on gelatin-coated slides, dehydrated in graded alcohol solutions, cleared in xylenes and coverslipped with Permount mounting media. As a negative control, a series of sections were processed without the primary antibody and did not produce any staining pattern.

#### StainAI

The StainAI model has previously been trained to detect six activation classes corresponding to ramified, hypertrophic, bushy, ameboid, rod-shaped, and hypertrophic rod-shaped microglia cells(*89*). Iba-1 immunostained sections were then scanned at 20x magnification with a resolution of 0.37 µm/pixel for StainAI analysis according to previously published methods(*89*). Briefly, A YOLO plus UNet pipeline was used detect and segment microglia. Each image underwent cell detection using YOLOv5 focusing on soma and process of Iba-1-stained microglia. Bounding boxes of 119 x 119 µm^2^ were superimposed on each cell for UNet segmentation to capture the cell body and processes. The UNet model detected cells and created corresponding cell masks. A C5.0 decision tree classifier was used to categorize key features based on 28 morphometric parameters (6 area-, 5 perimeter-, 6 span-length related parameters; 4 span-length ratios, 2 circularities and 3 fractal parameters; 2 parameters describing intercellular distancing properties of microglia) from each cell mask that were derived from the UNet segmentation. Each single-cell mask was color-coded according to the morphological phenotype of ramified (green), hypertrophic (yellow), bushy (orange), ameboid (red), rod-shaped (cyan), and hypertrophic rod-shaped (blue). Cells that were out of focus or smaller than 30 µm^2^ were grayed out and excluded from measurement. MATLAB was used to draw regions of interest to quantify microglia phenotypes from the YOLO+UNet pipeline(*89*).

#### Design-based stereology

Pyramidal neuronal populations of the CA 1-3 fields of the hippocampus were quantified using previously described sampling parameters(*90*). Briefly, a section sampling frequency of 1 in every 30 cresyl-violet sections was selected. Topography of subfields was achieved using a 4x objective and superimposed counting frames (1600 µm^3^) were generated using the MicroBrightField Stereoinvestigator program (Williston, VT, USA). A standard x-y grid was generated (CA1 500 x 500 µm; CA2 250 x 250 µm; CA3 350 x 350µm) and neurons were counted using a 100x objective (1.3 N.A.). To estimate the population of nestin-positive neurons, the granular/subgranular region of the dentate gyrus was delineated using a 10x objective, superimposed counting frames (1600 µm^3^) were generated using the MicroBrightField Stereoinvestigator program (Williston, VT, USA), a 100 x 100 µm x-y grid was generated and neurons were counted using a 100x objective (1.3 N.A.).

Myelin staining was achieved using a 0.2% gold chloride solution(*91*). Sections were removed from antigen preserve, washed in PBS, mounted on gelatin-coated slides and allowed to dry overnight. Sections were incubated with a 0.2% gold chloride solution for 2 hours at room temperature, washed in three exchanges of water, fixed for 5 minutes in a 2.5% solution of sodium thiosulphate and rinsed in running tap water for 30 minutes. Sections were then dehydrated in graded alcohol solutions (50-95%), cleared in xylenes and coverslipped with Permount. Images were captured using the Olympus CellSens program attached to an Olympus microscope with DP23 camera. Magnification of the sections was 200x, using a 20x objective. Images were then analyzed using ImageJ/Fuji software. For areas with fine fibers, images were converted to 8-bit and a threshold was applied to isolate the particles of interest. A region of interest (ROI) was selected. The "Fit Rectangle" tool was used to standardize the counting area, ensuring that each analyzed region was of the same size. Particle analysis was then performed using the "Analyze Particles" function. The primary focus for particle count was myelin density, for which the data accounted for. The particle outlines were generated, and the results, including particle counts, were displayed for each sample. For larger fiber bundles like the corpus callosum, fornix, and cingulum, an ROI was selected and mean density was measured. The embedded scale bar on each image was used to calibrate pixels to microns for the analyses.

### Tissue dissociation and single cell sorting

All dissociation steps were performed in the presence of Protector RNAse inhibitor (Sigma 03335402001), transcription inhibitor actinomycin D (Sigma-Aldrich; Cat#: A1410), and translation inhibitor anisomycin (Sigma-Aldrich; Cat#: A9789) based on recommendations in PMID: 35260865(*92*). Tissue was dissociated into a single cell suspension using previously published methods(*93*) (PMID: 26687838). Briefly, tissue was chopped using a scalpel and incubated in papain (20 units/mL) at 34 °C for 45 min before rinsing with a protease inhibitor solution (ovomucoid) and mechanical trituration. Dissociated cells were incubated with 50 nM calcein-AM and 4 uM ethidium homidimer-1 from the LIVE/DEAD™ Viability/Cytotoxicity Kit (Thermofisher Scientific, L3224) prior to cell sorting. Live (calcein+, ethidium-) cells were then sorted on a BD Facs Aria II. Cells were sorted into 4% BSA with RNAse inhibitor.

### Single cell RNA sequencing

Cell suspensions were loaded onto the 10X Genomics Chromium Controller using the Chromium NextGEM Single Cell 5’ Library & Gel Bead kit to capture individual cells and barcoded gel beads within droplets for reverse transcription(*94*). The libraries were prepared according to manufacturer instructions and sequenced on an Illumina NovaSeq 6000 with a paired-end 26x91 configuration targeting a depth of 50,000 reads per cell. Cell Ranger software was used to perform demultiplexing of cellular transcript data, and mapping and annotation of unique molecular identifiers (UMIs) and transcripts for downstream data analysis.

### Bioinformatics pipeline

Alignment, filtering, barcode counting and UMI counting were performed with Cell Ranger v.6.0.2 (10x Genomics) using a reference built from a composite of the Rhesus Mmul10 genome with Ensembl annotation v100 and the Zika virus strain PRVABC59 assembly (Accession KU501215.1). Single-cell RNA sequencing (scRNA-seq) data were analyzed using the Seurat package (v4.0.0) in R (v 2024.09.0+375). For quality control, cells with fewer than 500 detected features or with >20% of reads mapping to mitochondrial genes or <200 detected features per cell were excluded. After filtering, each dataset was normalized, and variable features were identified using the variance-stabilizing transformation (VST) method.

Metadata annotations for each Seurat object including animal ID, brain region (amygdala, hippocampus, and striatum), and treatment group (Zika or Control), with additional metadata added post hoc to ensure consistent labeling across samples. Following preprocessing, all individual datasets were integrated using Seurat’s integration workflow. The integrated dataset was scaled, and principal component analysis (PCA) was performed. Dimensionality reduction was conducted using Uniform Manifold Approximation and Projection (UMAP). Cell clustering was performed at multiple resolution parameters (0.1 to 0.5) to explore clustering granularity.

UMAP projections were visualized, and cell type identities were assigned based on canonical marker gene expression. A total of 105,421 control and 94,975 ZIKV-infected cells were successfully captured and profiled for transcriptomic analysis.

### Diffusion MRI Data Acquisition and Processing

Diffusion Weighted Imaging (DWI) scans were acquired on a 3T Siemens Trio scanner (Malvern, PA) at the ENPRC Imaging Center using an 8-channel phase array coil and following published protocols(*95*). Subjects were scanned supine under isoflurane anesthesia (0.8-1% isoflurane, inhalation) using a head holder with ear bars and a mouthpiece to secure the head and avoid motion artifacts. A vitamin E capsule was taped on the right temple to identify the right brain hemisphere. Animals were intubated, administered dextrose/NaCl (I.V.) for hydration, and placed on an MRI-compatible heating pad to maintain body temperature. Physiological measures were monitored and maintained throughout the scans following ENPRC and IACUC protocols. After the scan and complete recovery from anesthesia, subjects were returned to their home cages.

DWI scans were acquired with a single-shot double spin-echo diffusion-weighted EPI sequence (voxel size=1.0x1.0x1.0mm^3^, b:0, 1000 s/mm2, 128 directions).

Preprocessing of DWI images included automatic removal of artifact-rich images was done using DTIPrep(*96*); susceptibility-related distortion correction, eddy current, and motion correction using topup(*97*) and eddy_openmp tools(*98*). A weighted least-square estimation was employed to produce a diffusion tensor image for each subject. Multi-atlas templates were used to create masks for the subjects using deformable registration(*99*) and majority voting. DTI images were skull stripped, and a study-specific atlas was built using DTI Atlas Builder(*100*). Fifteen fiber tracts (Corpus Callosum dorsal segment 1 (cc1d), Corpus Callosum dorsal segment 2 (cc2d), Corpus Callosum dorsal segment 3 (cc3d), Genu of Corpus Callosum (ccg), Splenium of of Corpus Callosum (ccs), Cingulum left and right (cg_l, cg_r), Cortico-spinal left and right (cst_l, cst_r), Inferior longitudinal left and right (ilf_l, ilf_r), Uncinate left and right (uf_l, uf_r), Fornix left and right (fornix_l, fornix_r) were propagated into the study-specific atlas from high-resolution DTI rhesus macaque atlas. These propagated tracts were voxelized and used as seed maps for fiber tractography. Tractography was performed automatically using AutoTract(*101*), followed by a manual inspection stage to further clean the tract definitions. The processed tracts from study-specific atlas were then mapped back into each subject’s DTI image space using deformation fields calculated during the atlas building stage. Fractional anisotropy (FA), axial diffusivity (AD), mean diffusivity (MD), and radial diffusivity (RD) were sampled along the tract’s arc using DTI Atlas Fiber Analyzer(*100*). These process steps yielded diffusion profiles for each tract, and the mean FA, RD, AD, and MD per white matter tract. DTI characterizes the properties of water diffusion in brain tissue. The water diffusion is anisotropic in white matter regions with axons that are aligned and densely packed. The DTI scalars are widely used to describe the magnitude and direction of the water diffusivity in the brain. FA is a measure of directional variation in diffusivities and is sensitive to changes in white matter micro-organization, but not specific to the type of the changes. FA should be interpreted along with the radial diffusivity RD and the axial diffusivity AD for more specific interpretations. RD reflects the magnitude of diffusion perpendicular to white matter fibers and sensitive to myelination level, loss of neurons, and reduction of axonal density. AD describes diffusion parallel to the fibers and is sensitive to axon loss or injury. Mean diffusivity MD increases in response to edema and cell loss and decreases with cell proliferation. The mean FA, AD, MD, and RD per tract were computed for the final analyses.

### Head Coil Change Adjustment

During data acquisition, the MRI head coil became inoperative and was replaced with the same model of coil. To assess the effect of the coil change, we compared the results before and after the coil swap. The new coil led to the significant increase in FA values. Given the small sample size, we adjusted for this coil effect using the mean shift method instead of using coil as additional covariate in the analysis. Specifically, we randomly selected two male subjects and two female subjects from both the ZIKV and Control groups scanned before and after the coil change (resulting in six scans before the coil change and six scans after). For each white matter fiber tract, we calculated the mean values of FA, AD, RD, and MD obtained using older and newer coils. Then we computed the differences in these values between scans taken with the old and new coils and subtracted from the data obtained using newer coil.

### Statistical Analyses

To ensure that infants randomly assigned to each group did not differ at the beginning of the experiment, physical measurements were analyzed using an independent T-test and Cohen’s D effect size. To assess potential neurobehavioral differences, the four INAS scores were analyzed separately used a Linear Mixed Model with Group (ZIKV-infected, uninfected control) and Age (3-6 weeks) as fixed factors and individual animal as a random factor. Results were graphed using GraphPad Prism 10.01 (GraphPad Software) and behavioral data was analyzed using SPSS 29 for Windows (IBM), significance was set at p < 0.05, and effect sizes were calculated using ηp^2^.

Differential gene expression (DGE) analysis between ZIKV-infected and control animals was performed using the FindMarkers function in Seurat, which employs the Wilcoxon Rank Sum test. Bonferroni-adjusted *p*-values were reported to account for multiple testing. Top differentially expressed genes were additionally filtered based on a Bonferroni-adjusted *p*-value threshold of < 0.05. Gene Ontology (GO) term enrichment analysis was subsequently performed on the filtered gene sets using the g:Profiler web tool (https://biit.cs.ut.ee/gprofiler/gost). Gene Set Enrichment Analysis (GSEA) was performed using the fgsea R package (v1.30.0) to assess the enrichment of curated custom gene sets among transcriptional changes of various cell types between ZIKV-infected and control animals. Differential expression results generated from Seurat were used to create a ranked gene list based on average log2fold-change values.

Enrichment was evaluated by comparing the ranked gene list against the custom gene set using fgsea, with 1,000 permutations and a fixed seed for reproducibility. Enrichment statistics including the Normalized Enrichment Score (NES), Enrichment Score (ES), and nominal *p*-value were reported.

Data were analyzed using a series of two-way analyses of variance (ANOVA) with Group (Zika vs. Control) and Sex (Male vs. Female) as between-subjects factors. For each tract/region, ANOVA models were fitted to test for main effects of Group and Sex. Where appropriate, Tukey’s post hoc multiple comparisons were conducted to further examine group differences while controlling the family-wise error rate. Model assumptions were assessed by inspecting residuals and formally testing the homogeneity of variances using Levene’s test. For each effect, sums of squares, degrees of freedom, F-statistics, and p-values were reported.

## Data and code availability

- All sequencing data will be deposited in the Gene Expression Omnibus (GEO) repository and accession provided at time of publication.
- Histology, behavior and MRI data reported in this paper will be shared by the lead contact upon request.
- Any additional information required to reanalyze the data reported in this paper is available from the lead contact upon request.

## Author contributions

Conceptualization: S.A.S., J.R., and A.C. Investigation:

- Animal care, infection, neurobehavioral assessment and brain dissection: S.F., R.R., and J.R.
- ZIKA RNA detection by PCR: N.S., K.M.M., and M.S.S.
- Histological analysis: A.H., Y.M.A., S.A., E.N., A.M., C.H.H., T.W.T., and M.B.
- Tissue dissociation and single cell sorting: M.M.S^1^, and S.A.S.
- Single cell RNA sequencing and bioinformatics pipeline: M.J.L., G.K.T., and S.E.B.
- Diffusion MRI data acquisition and processing: J.R., M.S., and R.V.
- Statistical analysis: V.V.E., R.V., J.R., and A.C.

Writing – original draft: V.V.E., S.A.S., R.V., M.B., M.J.L., G.K.T., M.M.S^2^., J.R., and A.C.

Writing – review and editing: V.V.E., S.A.S., R.V., M.B., M.M.S^1^, K.M.M., M.M.S^2^., J.R., and A.C.

^1^M.M.S: Maureen M. Sampson

^2^M.M.S: Mar M. Sanchez

## Acknowledgements

We would like to thank the Emory National Primate Research Center Veterinarians and Division of Research Resources staff for their expert assistance.

## Materials & Correspondence

### Lead contact

Further information and requests for resources and reagents should be directed to, and will be fulfilled by, the lead contact, Ann Chahroudi.

## Materials availability

All unique reagents, including available NHP samples, generated from this study are available from the lead contact with a completed materials transfer agreement.

---

**Supplemental Figure 1.**
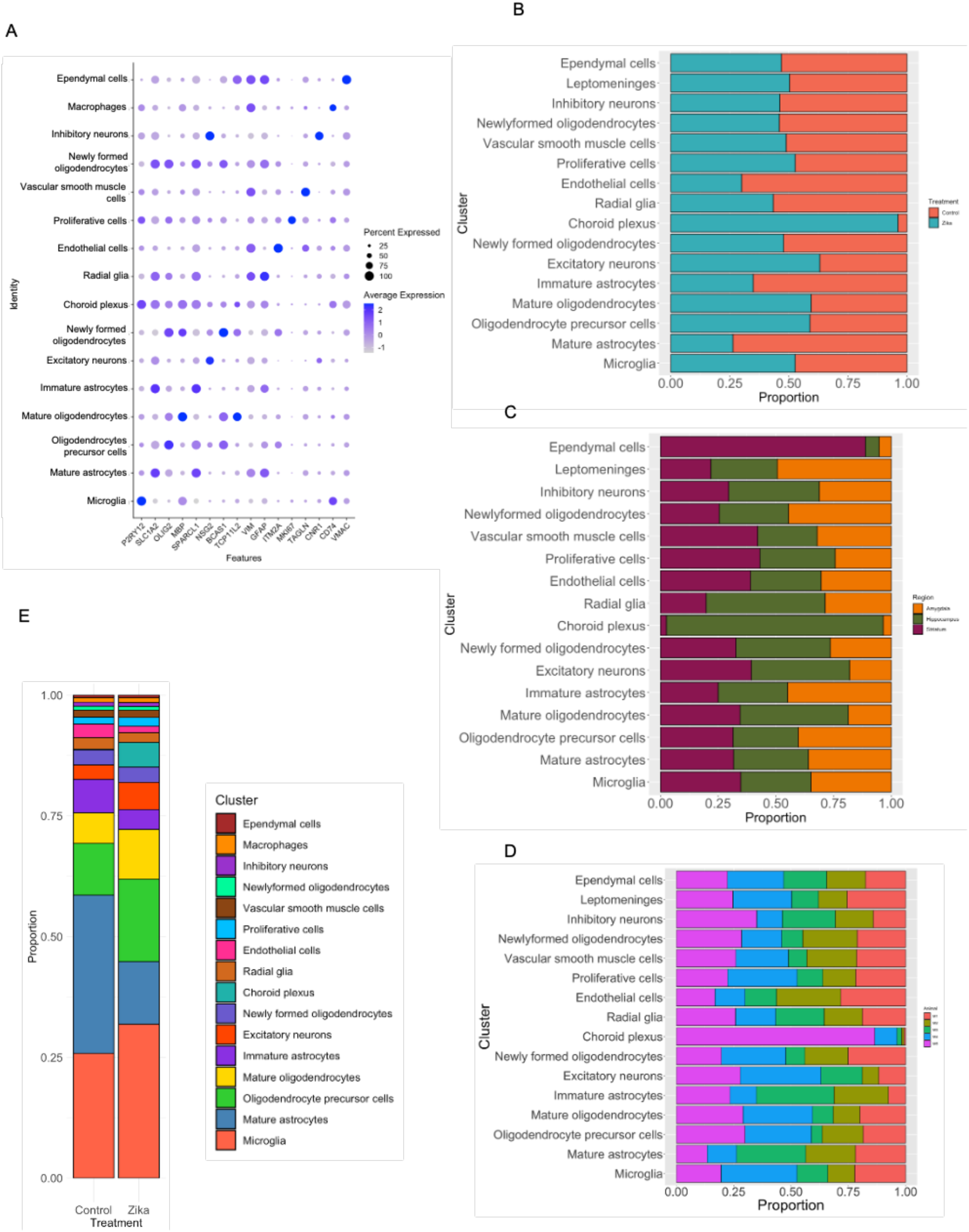
Characterization and distribution of cell type clusters across treatment groups, brain regions and individual animal, related to Figure 1. **(A)** Cell type identities were assigned based on canonical marker gene expression. Dot plot showing scaled expression and percent of cells expressing representative marker genes for each identified cell type cluster. **(B-D)** Bar plots showing proportion of cells within each cluster, stratified by treatment group **(B)**, brain region **(C)**, and individual animal **(D**). Bars represent the relative abundance of each cluster within the specified category. **(E)** Bar plot showing the relative proportions of all identified cell type clusters across control and ZIKV-infected groups.

**Supplemental Figure 2.**
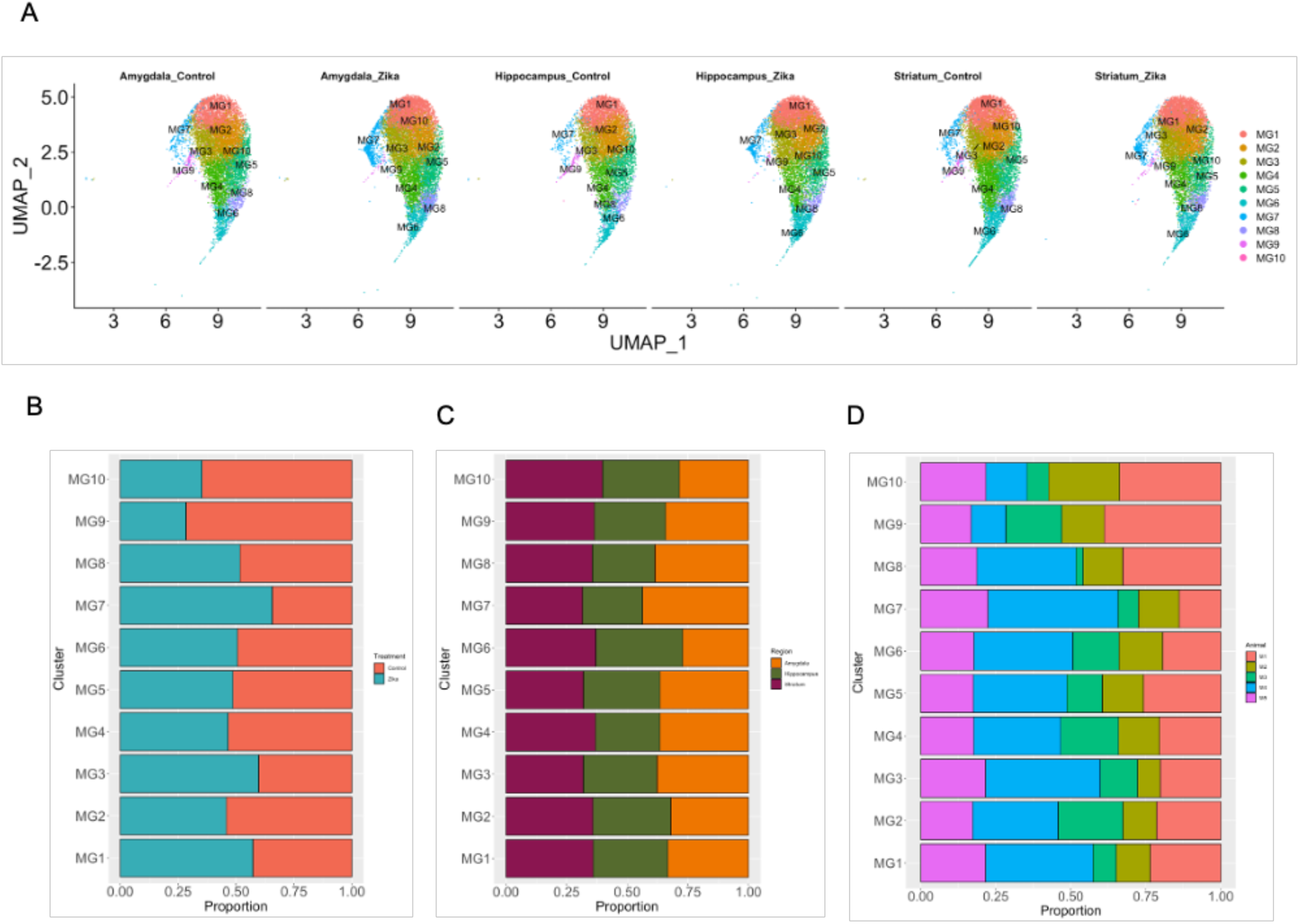
UMAP visualization and distribution of microglial subclusters across treatment groups, brain regions and individual animal, related to Figure 2. **(A)** UMAP projections of microglia subclusters from the amygdala, hippocampus, and striatum, with each region’s cells split by treatment group (ZIKV-infected vs. control). Subclusters were identified based on transcriptional profiles and are color-coded consistently across plots. **(B-D)** Bar plots showing the proportion of each microglial subcluster across treatment groups **(B)**, brain regions **(C)**, and individual animal **(D)**.

**Supplemental Figure 3.**
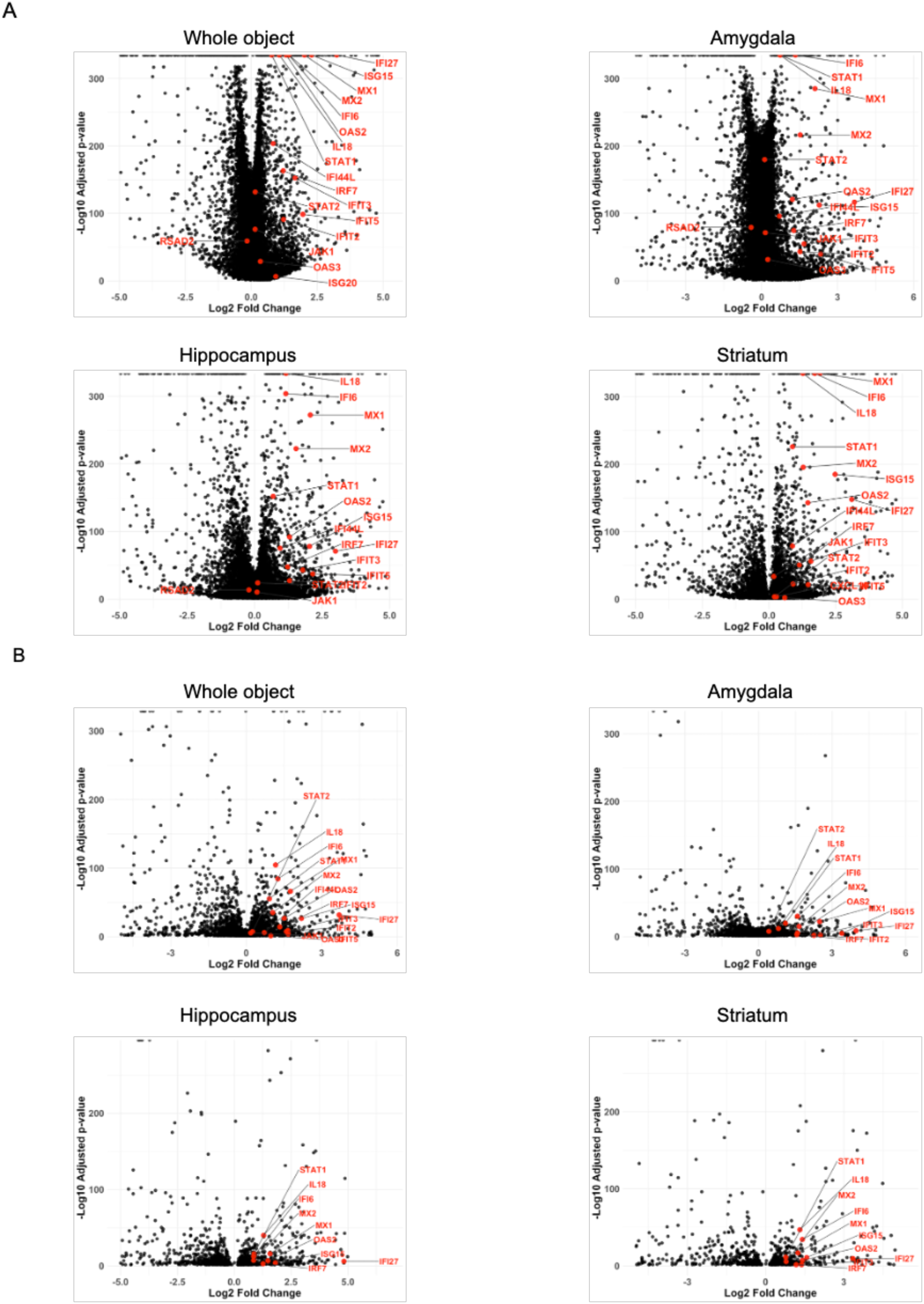
Gene expression in total microglia and MG6 subcluster across brain regions shows induction of interferon response genes, related to Figure 2. **(A)** Volcano plots showing differential expression of a curated panel of interferon-stimulated genes (ISGs) in the total microglial population, comparing ZIKV-infected and control infant rhesus macaques in the combined dataset and within individual brain regions (amygdala, hippocampus, and striatum). **(B)** Volcano plots of ISG expression within the MG6 subcluster, analyzed similarly across the combined dataset and individual brain regions. Each black point represents an individual gene, with interferon-stimulated genes (ISGs) highlighted in red. Plots display log₂ fold change on the x-axis and –log₁₀(p-value) on the y-axis.

**Supplemental Figure 4.**
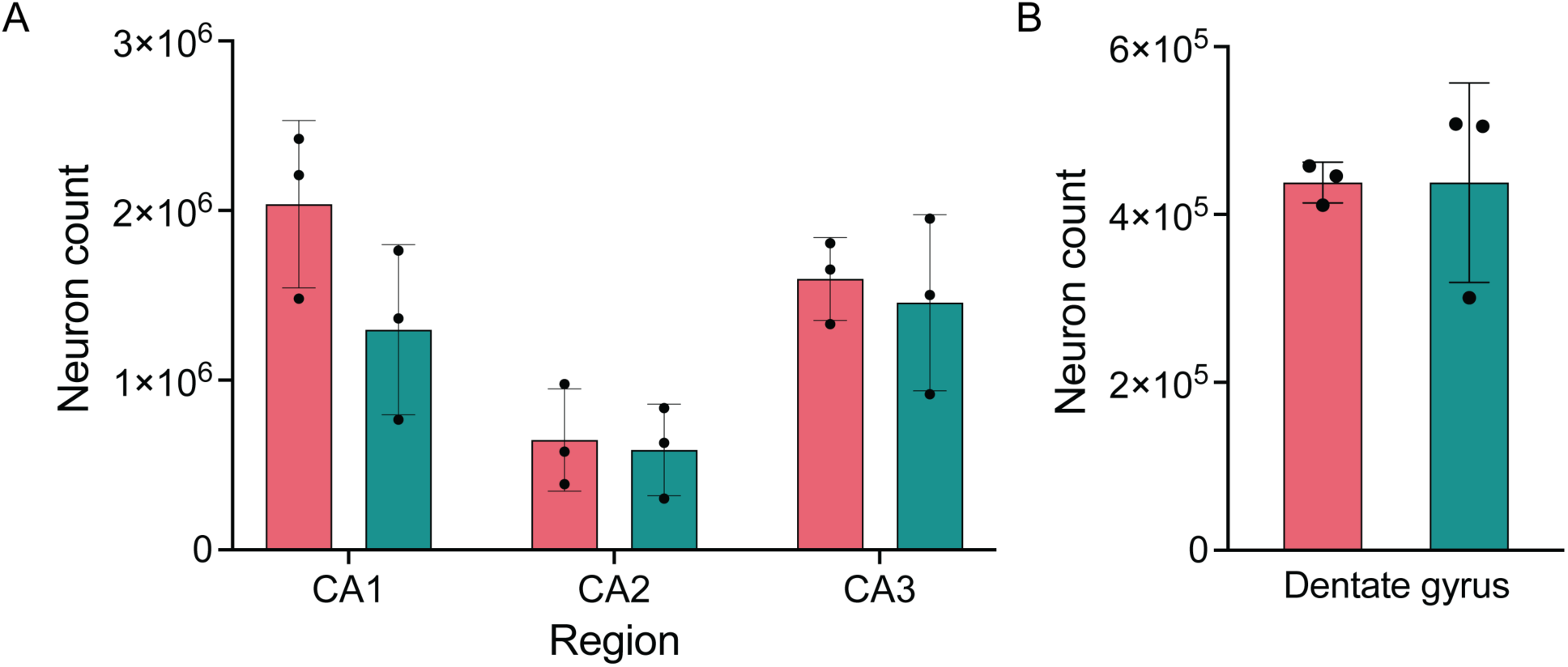
Zika virus infection induce neuronal apoptosis in the hippocampal CA1 region, related to Figure 3. **(A)** Design-based stereologic quantification of hippocampal neuronal populations of the pyramidal layer in CA subfields 1-3 and **(B)** Design-based stereologic quantification of nestin positive immature neurons in the dentate gyrus.

**Supplemental Figure 5.**
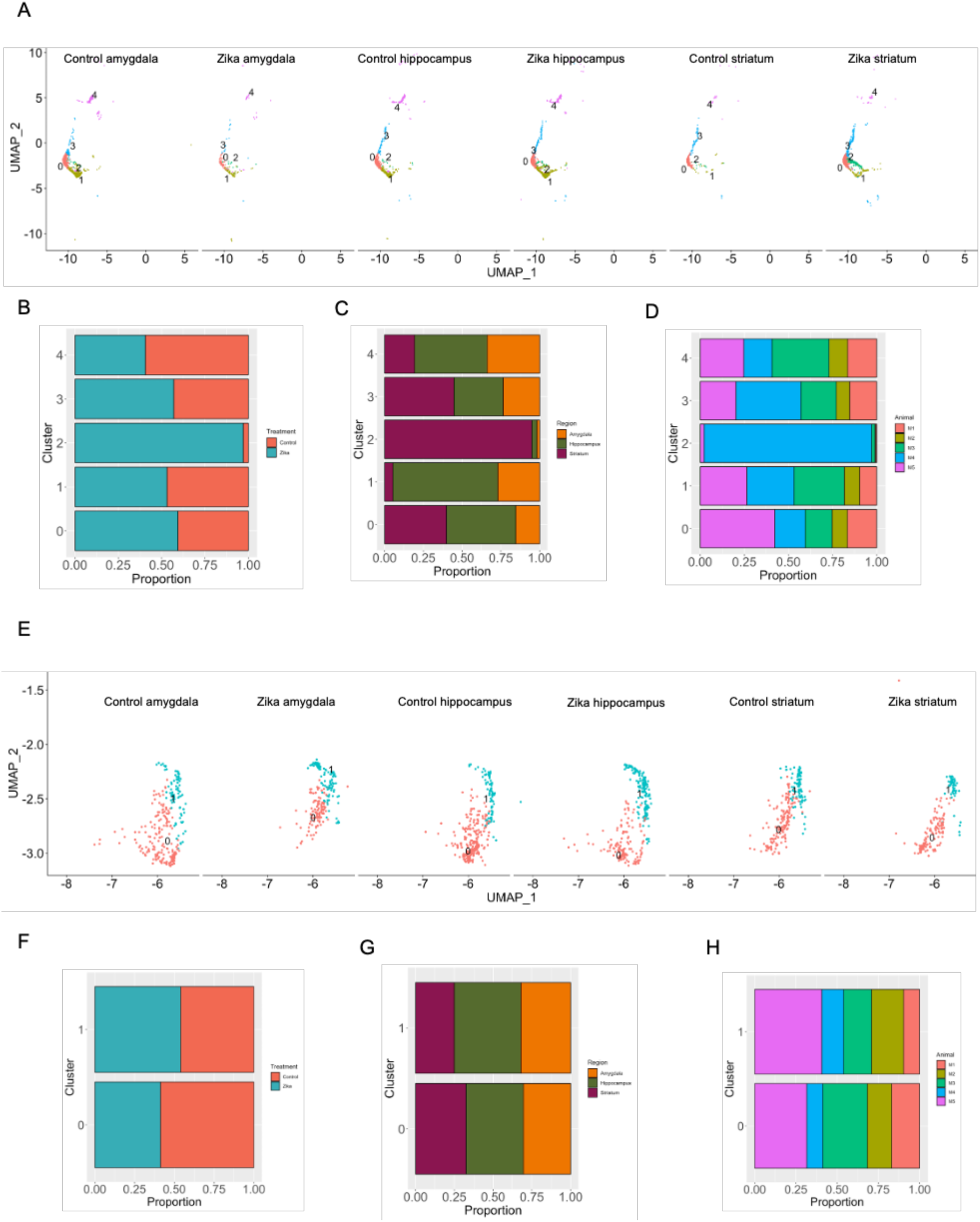
UMAP visualization and distribution of excitatory and inhibitory neuron subclusters across treatment groups, brain regions and individual animal, related to Figure 3. **(A)** UMAP projections of excitatory neuron subclusters from the amygdala, hippocampus, and striatum, with each region’s cells split by treatment group (ZIKV-infected vs. control). Subclusters are color-coded consistently across plots. **(B-D)** Bar plots showing the proportion of each excitatory neuron subcluster across treatment groups **(B)**, brain regions **(C)**, and individual animal **(D)**. **(E)** UMAP projections of inhibitory neuron subclusters from the amygdala, hippocampus, and striatum, with each region’s cells split by treatment group (ZIKV-infected vs. control). Subclusters are color-coded consistently across plots. **(F-H)** Bar plots showing the proportion of each inhibitory neuron subcluster across treatment groups **(F)**, brain regions **(G)**, and individual animal **(H)**.

**Supplemental Figure 6.**
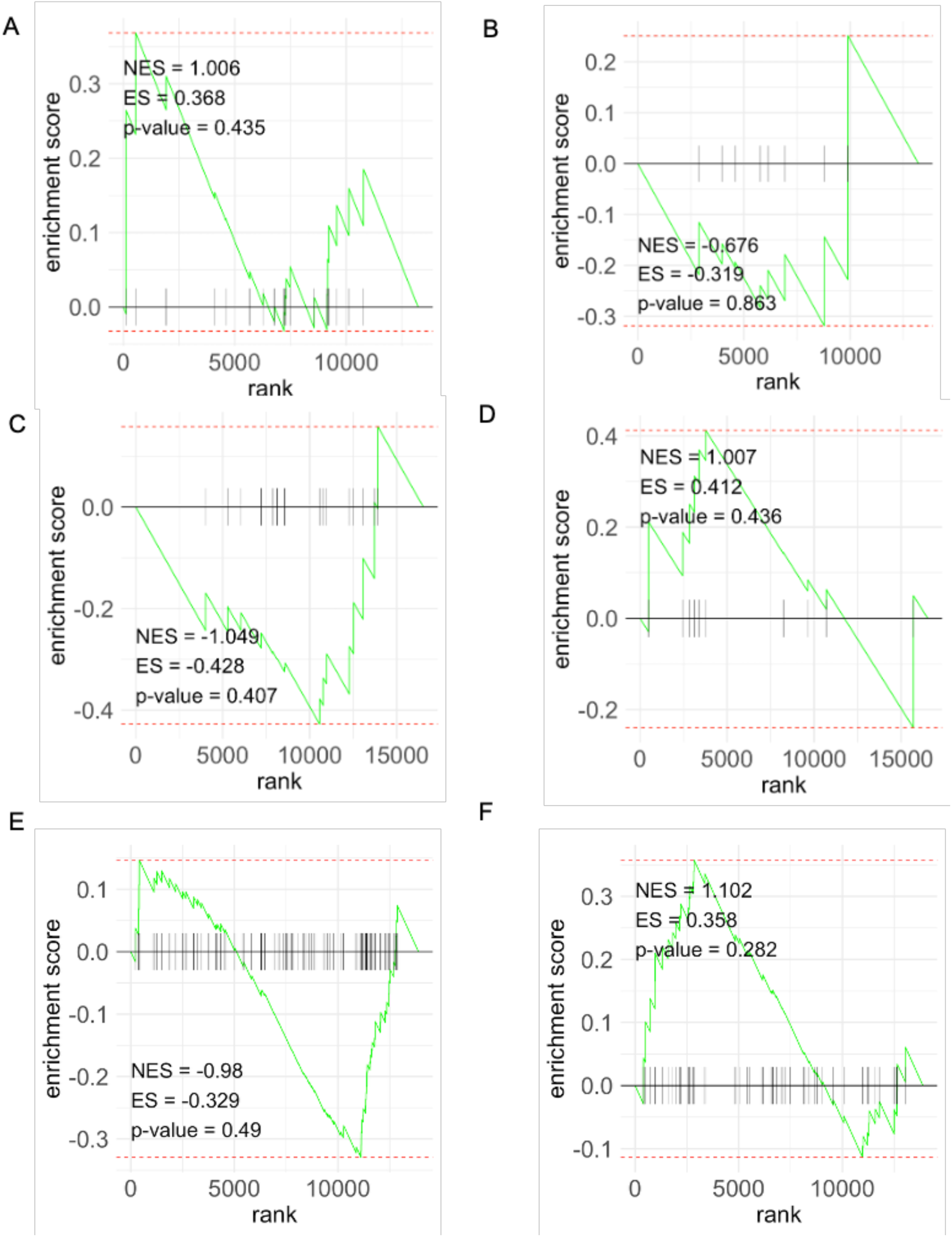
Gene Set Enrichment Analysis (GSEA) of cerebral palsy and autism spectrum disorder (ASD) gene sets in neuronal and mature oligodendrocyte populations show selective enrichment of ASD genes in neuronal populations, related to Figure 4. **(A & B)** GSEA plots of cerebral palsy-associated gene sets in excitatory neurons, derived from the Clinical Genome Resources **(A)** and Harmonizome **(B)** databases. **(C & D)** GSEA plots of cerebral palsy-associated gene sets in inhibitory neurons, derived from the Clinical Genome Resources **(C)** and Harmonizome **(D)** databases. **(E & F)** GSEA plots of Epoch 1 **(E)** and Epoch 2 **(F)** ASD gene sets in mature oligodendrocytes. Enrichment scores and significance values reflect the degree of overrepresentation of disease-associated gene sets within the specified cell populations.

**Supplemental Figure 7.**
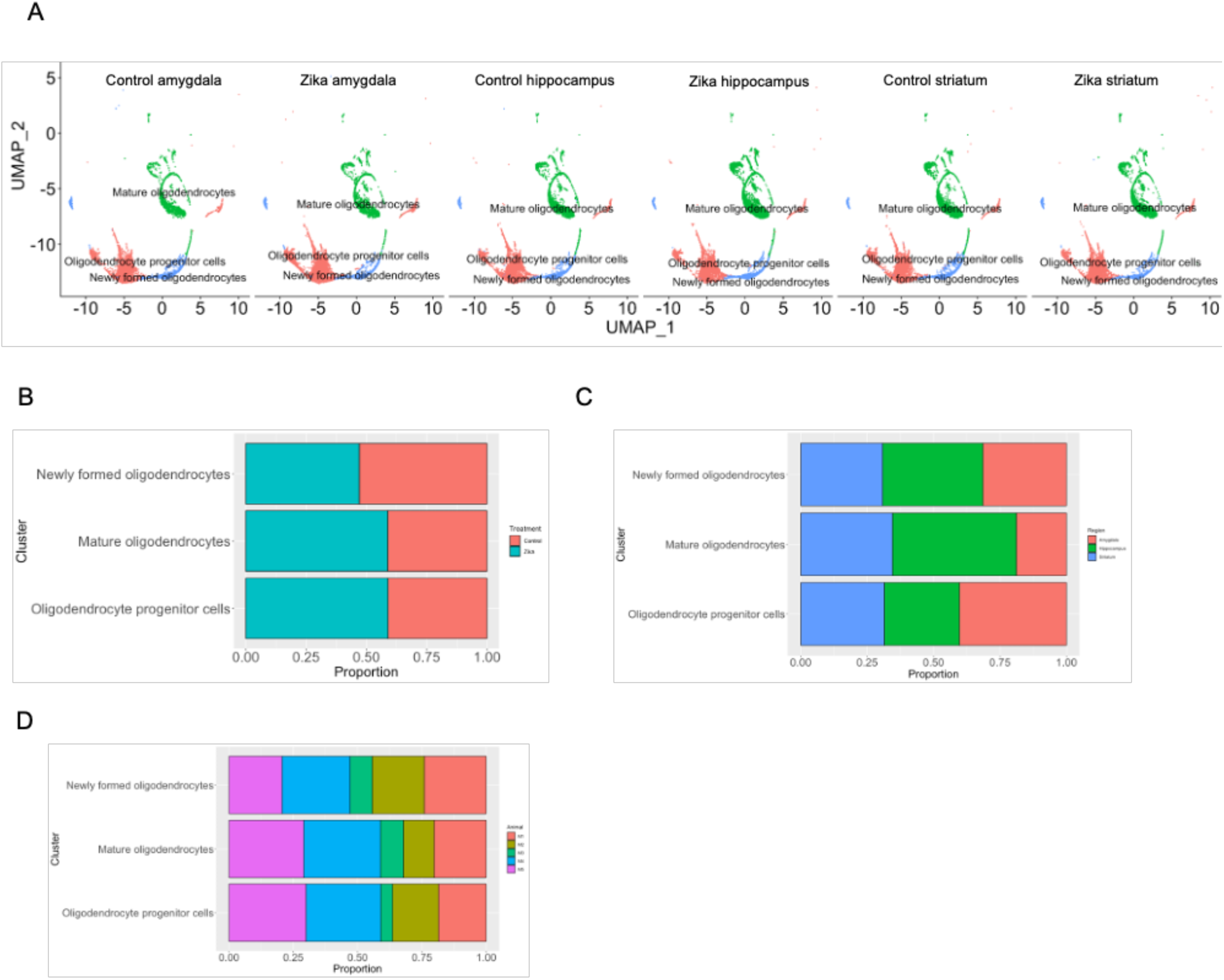
UMAP visualization and distribution of oligodendrocyte subclusters related to Figure 5. **(A)** UMAP projections of oligodendrocyte subclusters from the amygdala, hippocampus, and striatum, with each region’s cells split by treatment group (ZIKV-infected vs. control). Subclusters were identified based on transcriptional profiles and are color-coded consistently across plots. **(B-D)** Bar plots showing the proportion of each oligodendrocyte subcluster across treatment groups **(B)**, brain regions **(C)**, and individual animal **(D)**.

**Supplemental Figure 8.**
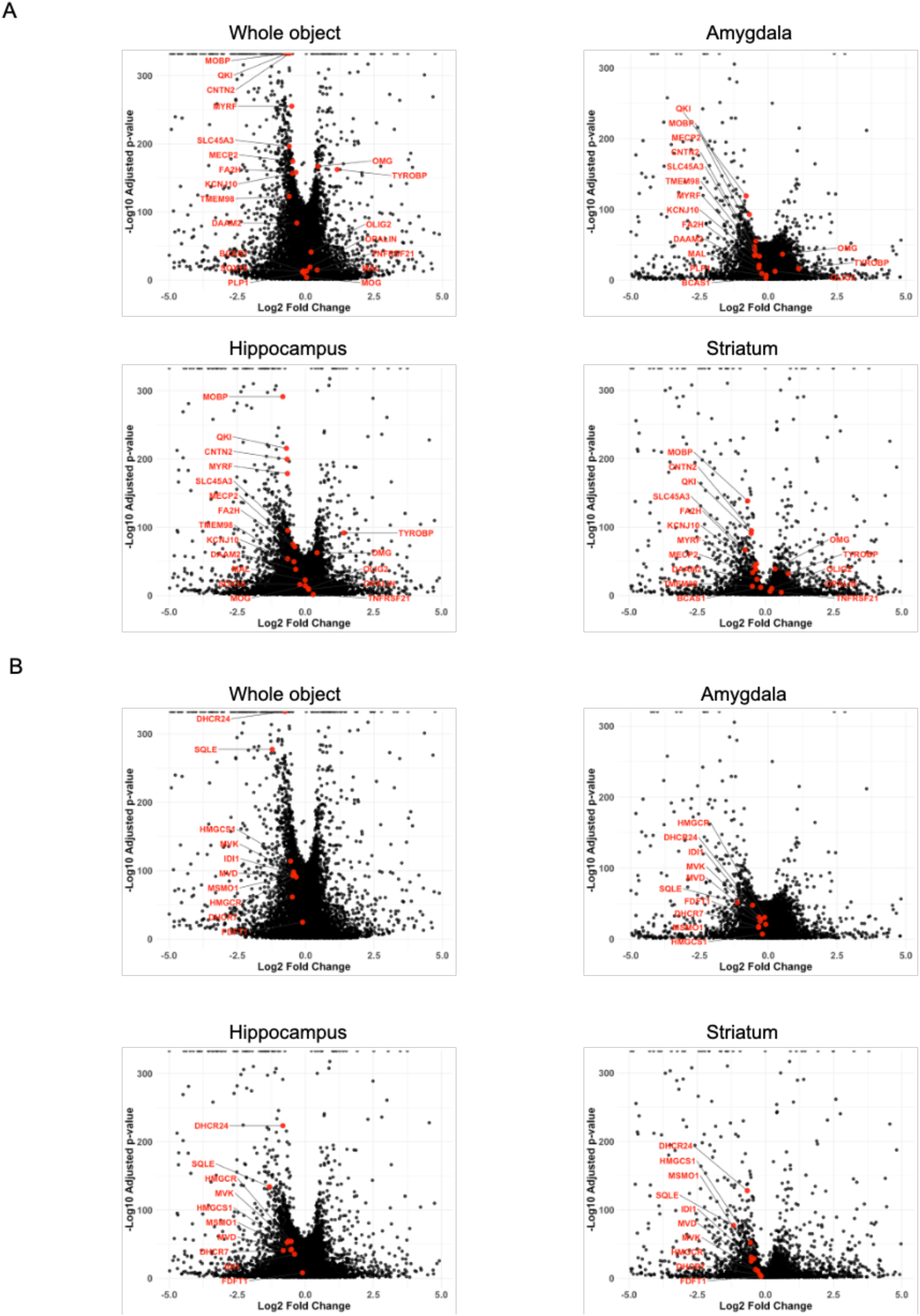
Gene expression in mature oligodendrocytes across brain regions shows downregulation of representative genes from sterol biosynthesis and oligodendrocyte differentiation pathways, related to Figure 5. **(A)** Volcano plot illustrating differential expression of key genes involved in the sterol biosynthesis pathway between experimental groups in the combined dataset and within individual brain regions (amygdala, hippocampus, and striatum). **(B)** Volcano plot showing differential expression of genes associated with oligodendrocyte differentiation across the combined dataset and individual brain regions. Each point represents an individual gene, plotted by log₂ fold change (x-axis) versus –log₁₀(p-value) (y-axis). Genes highlighted in red color are representative members of the respective pathways.

**Supplemental Figure 9.**
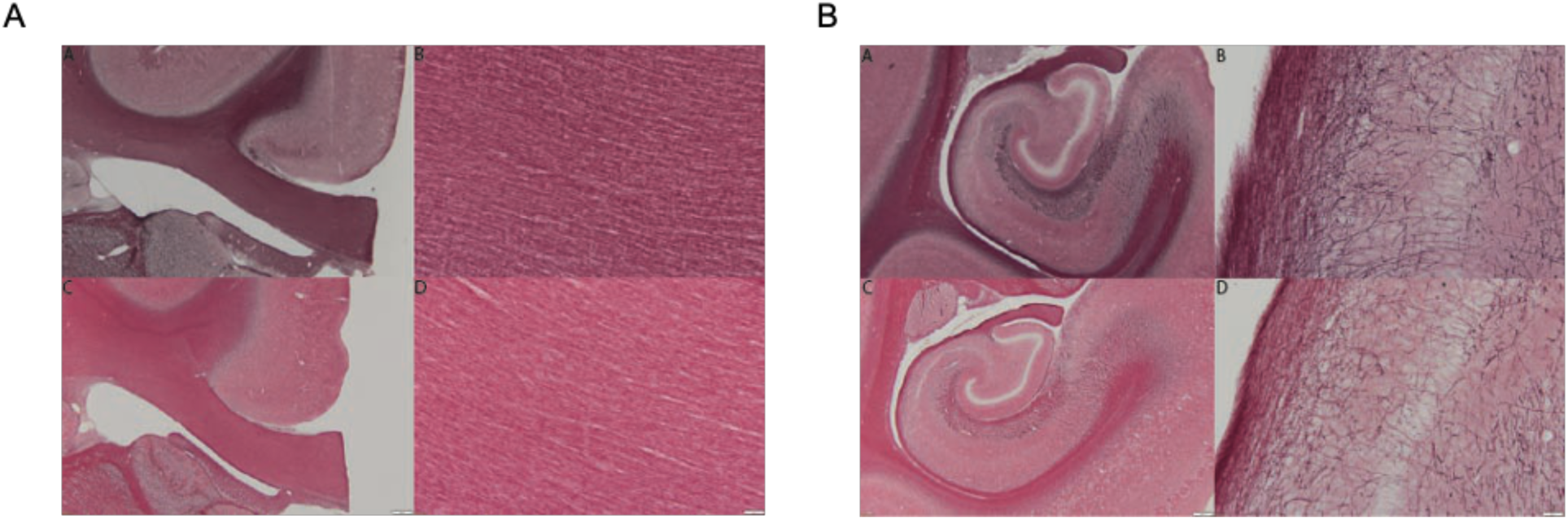
Zika virus infection did not alter myelin density in corpus callosum and limbic system white matter tracts, related to Figure 6. Matched sections from each subject were stained with a 0.2% gold chloride solution from **(A)** corpus callosum and **(B)** hippocampal region. Density analysis with Image J/Fuji did not reveal significant differences in myelin density between groups in either area. Top panels are representative images from control and bottom panels are from Zika-infected subjects (scale bar left panels=200 µm; scale bars right panels=20 µm).

**Supplementary Table 1.**
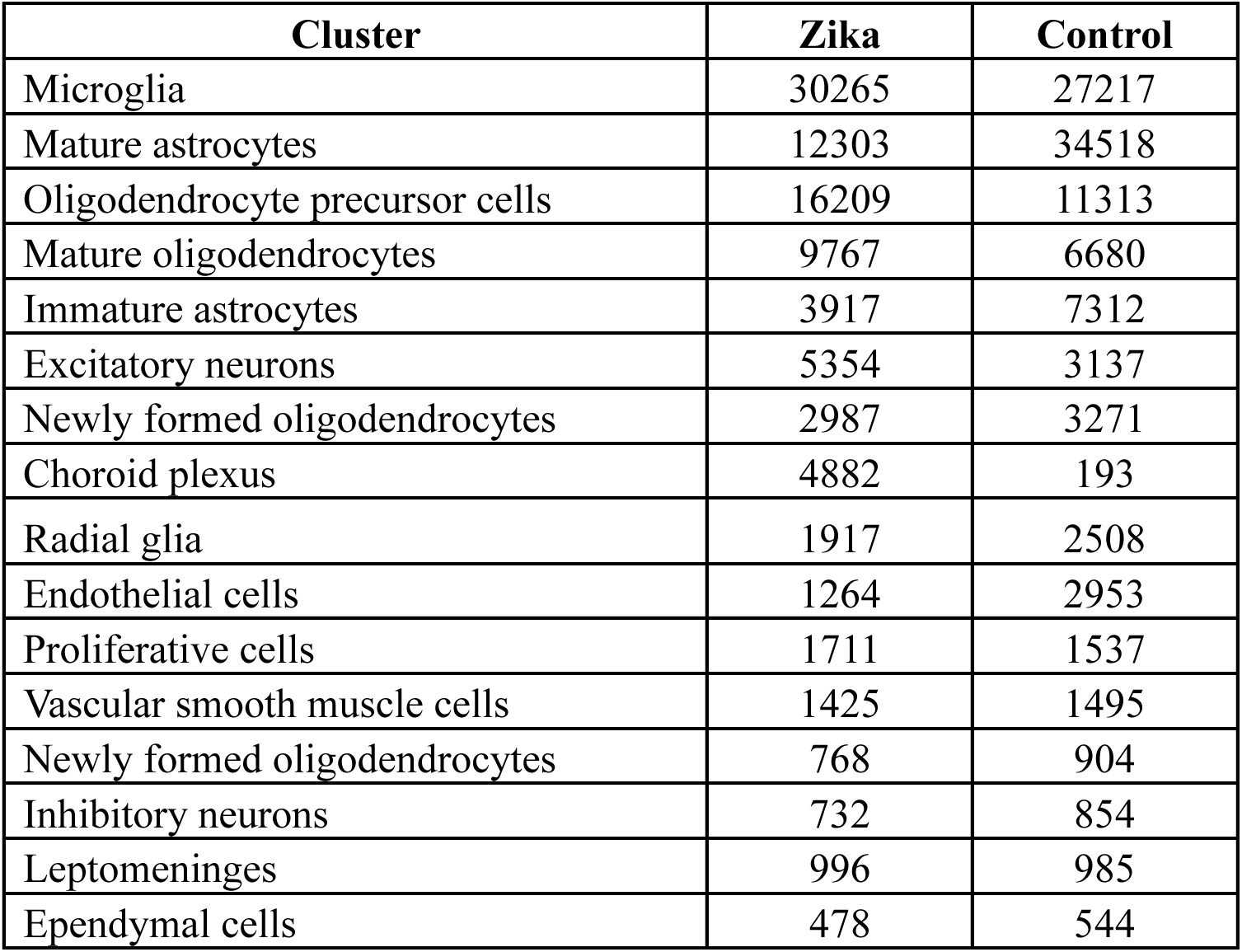
Cell numbers for all 16 clusters by experimental group.

**Supplementary Table 2.**
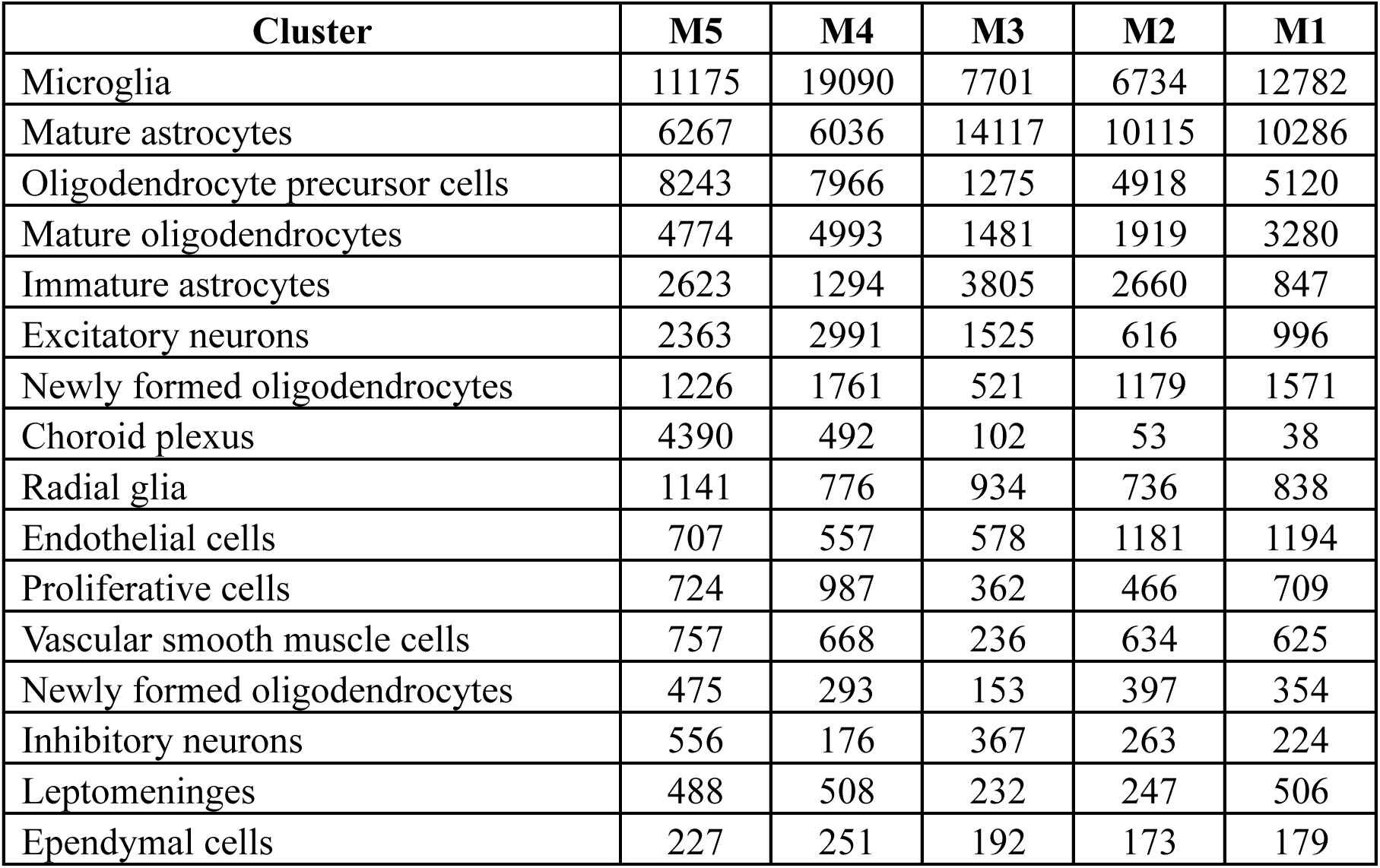
Cell numbers for all 16 clusters by individual animal.

| **Cluster** | **M5** | **M4** | **M3** | **M2** | **M1** |
| --- | --- | --- | --- | --- | --- |
| Microglia | 11175 | 19090 | 7701 | 6734 | 12782 |
| Mature astrocytes | 6267 | 6036 | 14117 | 10115 | 10286 |
| Oligodendrocyte precursor cells | 8243 | 7966 | 1275 | 4918 | 5120 |
| Mature oligodendrocytes | 4774 | 4993 | 1481 | 1919 | 3280 |
| Immature astrocytes | 2623 | 1294 | 3805 | 2660 | 847 |
| Excitatory neurons | 2363 | 2991 | 1525 | 616 | 996 |
| Newly formed oligodendrocytes | 1226 | 1761 | 521 | 1179 | 1571 |
| Choroid plexus | 4390 | 492 | 102 | 53 | 38 |
| Radial glia | 1141 | 776 | 934 | 736 | 838 |
| Endothelial cells | 707 | 557 | 578 | 1181 | 1194 |
| Proliferative cells | 724 | 987 | 362 | 466 | 709 |
| Vascular smooth muscle cells | 757 | 668 | 236 | 634 | 625 |
| Newly formed oligodendrocytes | 475 | 293 | 153 | 397 | 354 |
| Inhibitory neurons | 556 | 176 | 367 | 263 | 224 |
| Leptomeninges | 488 | 508 | 232 | 247 | 506 |
| Ependymal cells | 227 | 251 | 192 | 173 | 179 |

**Supplementary Table 3.** Cell numbers for all 16 clusters by brain region.

| **Cluster** | **Striatum** | **Hippocampus** | **Amygdala** |
| --- | --- | --- | --- |
| Microglia | 19983 | 17511 | 19988 |
| Mature astrocytes | 14847 | 15163 | 16811 |
| Oligodendrocyte precursor cells | 8660 | 7768 | 11094 |
| Mature oligodendrocytes | 5672 | 7719 | 3056 |
| Immature astrocytes | 2806 | 3375 | 5048 |
| Excitatory neurons | 3341 | 3618 | 1532 |
| Newly formed oligodendrocytes | 2048 | 2557 | 1653 |
| Choroid plexus | 131 | 4766 | 178 |
| Radial glia | 876 | 2279 | 1270 |
| Endothelial cells | 1645 | 1289 | 1283 |
| Proliferative cells | 1401 | 1057 | 790 |
| Vascular smooth muscle cells | 1229 | 752 | 939 |
| Newly formed oligodendrocytes | 426 | 501 | 745 |
| Inhibitory neurons | 469 | 622 | 495 |
| Leptomeninges | 432 | 570 | 979 |
| Ependymal cells | 907 | 61 | 54 |

**Supplementary Table 4.**
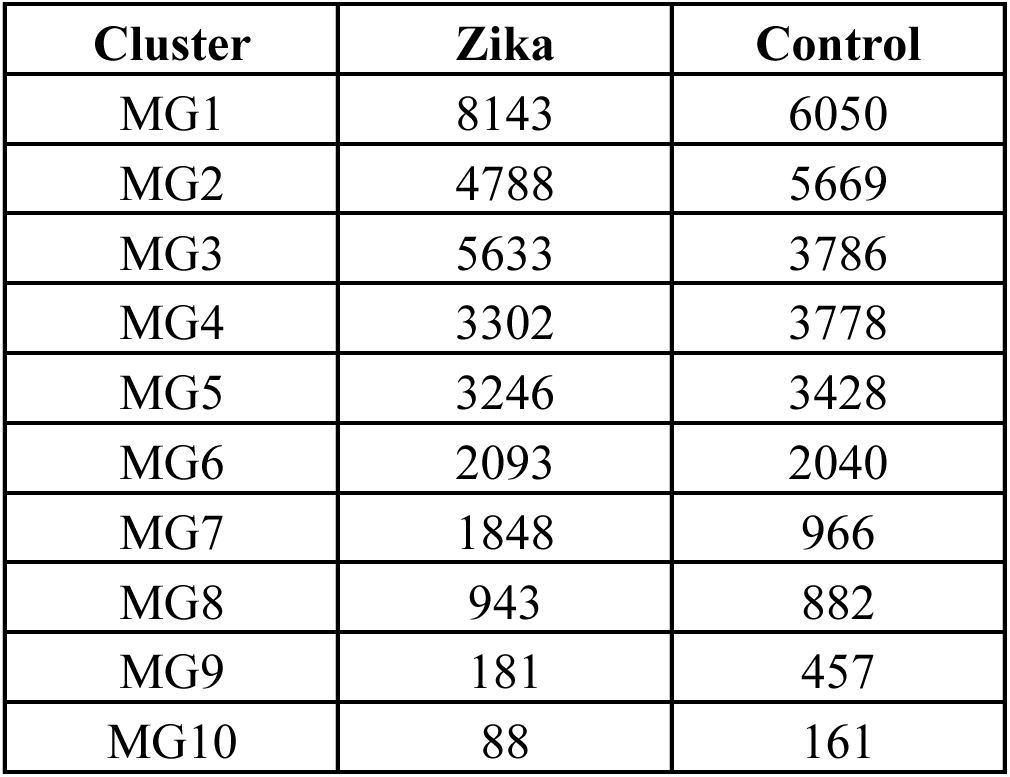
Cell numbers for all microglial subclusters by experimental group.

**Supplementary Table 5.** Cell numbers for all microglial subclusters by individual animal.

| **Cluster** | **M5** | **M4** | **M3** | **M2** | **M1** |
| --- | --- | --- | --- | --- | --- |
| MG1 | 3063 | 5080 | 1101 | 1608 | 3341 |
| MG2 | 1804 | 2984 | 2255 | 1173 | 2241 |
| MG3 | 2030 | 3603 | 1175 | 703 | 1908 |
| MG4 | 1246 | 2056 | 1366 | 963 | 1449 |
| MG5 | 1170 | 2076 | 792 | 916 | 1720 |
| MG6 | 730 | 1363 | 637 | 601 | 802 |
| MG7 | 630 | 1218 | 193 | 381 | 392 |
| MG8 | 341 | 602 | 45 | 240 | 597 |
| MG9 | 107 | 74 | 119 | 90 | 248 |
| MG10 | 54 | 34 | 18 | 59 | 84 |

**Supplementary Table 6.**
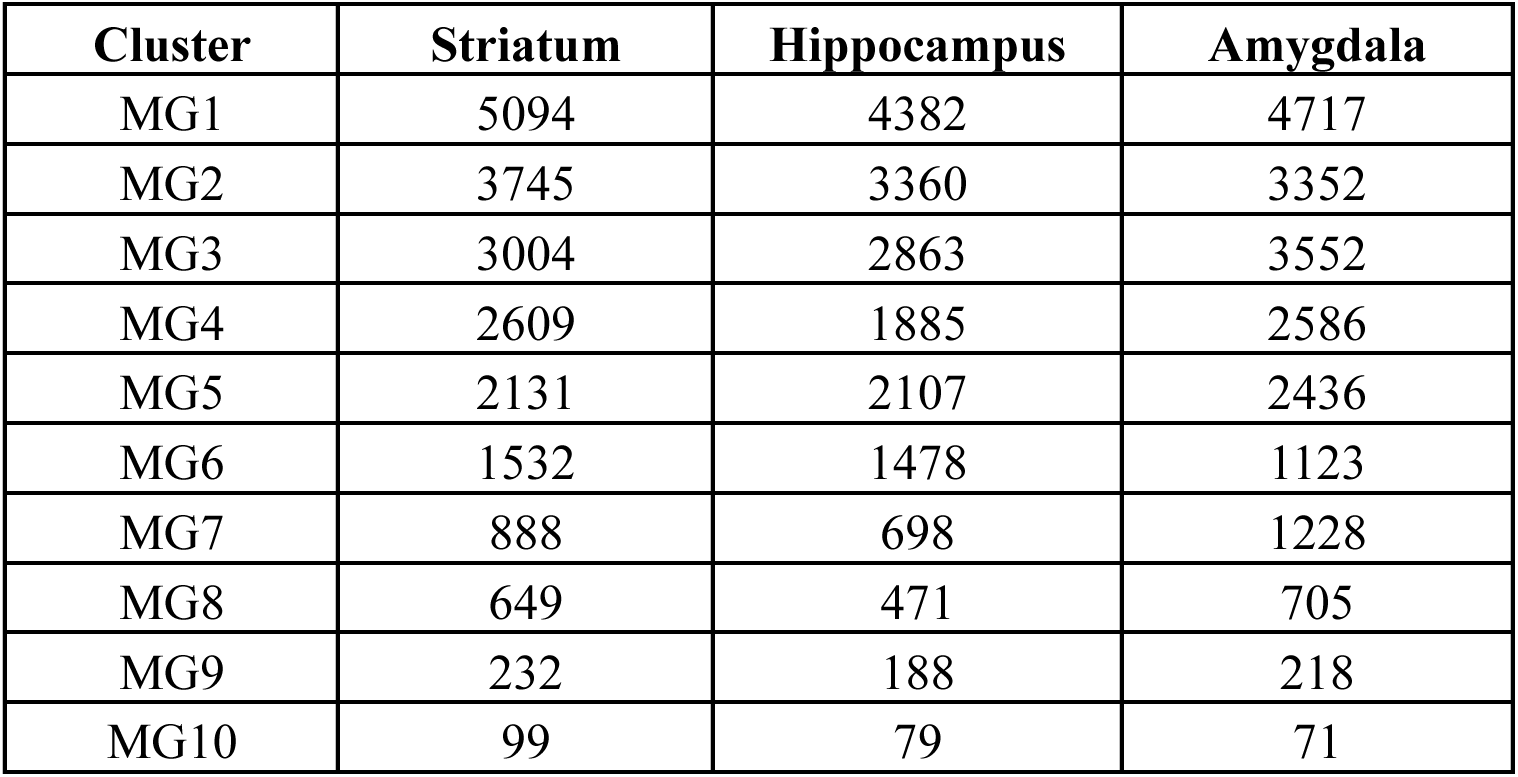
Cell numbers for all microglial subclusters by brain region.

**Supplementary Table 7.** Cell numbers for all excitatory neuron subclusters by experimental group.

| **Cluster** | **Zika** | **Control** |
| --- | --- | --- |
| 0 | 2018 | 1379 |
| 1 | 1247 | 1101 |
| 2 | 1459 | 44 |
| 3 | 436 | 329 |
| 4 | 194 | 284 |

**Supplementary Table 8.** Cell numbers for all excitatory neuron subclusters by individual animal.

| Cluster | M5 | M4 | M3 | M2 | M1 |
| --- | --- | --- | --- | --- | --- |
| 0 | 1436 | 582 | 517 | 298 | 564 |
| 1 | 622 | 625 | 670 | 204 | 227 |
| 2 | 32 | 1427 | 29 | 6 | 9 |
| 3 | 155 | 281 | 154 | 58 | 117 |
| 4 | 118 | 76 | 155 | 50 | 79 |

**Supplementary Table 9.** Cell numbers for all excitatory neuron subclusters by brain region.

| Cluster | Striatum | Hippocampus | Amygdala |
| --- | --- | --- | --- |
| 0 | 1356 | 1516 | 525 |
| 1 | 124 | 1588 | 636 |
| 2 | 1427 | 48 | 28 |
| 3 | 342 | 242 | 181 |
| 4 | 92 | 224 | 162 |

**Supplementary Table 10.** Cell numbers for all inhibitory neuron subclusters by experimental group.

| Cluster | Zika | Control |
| --- | --- | --- |
| 0 | 399 | 569 |
| 1 | 333 | 285 |

**Supplementary Table 11.** Cell numbers for all inhibitory neuron subclusters by individual animal.

| Cluster | M5 | M4 | M3 | M2 | M1 |
| --- | --- | --- | --- | --- | --- |
| 0 | 305 | 94 | 262 | 143 | 164 |
| 1 | 251 | 82 | 105 | 120 | 60 |

**Supplementary Table 12.** Cell numbers for all inhibitory neuron subclusters by brain region.

| Cluster | Striatum | Hippocampus | Amygdala |
| --- | --- | --- | --- |
| 0 | 315 | 357 | 296 |
| 1 | 154 | 265 | 199 |

**Supplementary Table 13.** Cell numbers for all oligodendrocyte subclusters by experimental group.

| **Cluster** | **Zika** | **Control** |
| --- | --- | --- |
| Oligodendrocyte progenitor cells | 16174 | 11255 |
| Mature oligodendrocytes | 10189 | 7122 |
| Newly formed oligodendrocytes | 3368 | 3791 |

**Supplementary Table 14.** Cell numbers for all oligodendrocyte subclusters by individual animal.

| **Cluster** | **M5** | **M4** | **M3** | **M2** | **M1** |
| --- | --- | --- | --- | --- | --- |
| Oligodendrocyte progenitor cells | 8218 | 7956 | 1267 | 4893 | 5095 |
| Mature oligodendrocytes | 5026 | 5163 | 1545 | 2072 | 3505 |
| Newly formed oligodendrocytes | 1474 | 1894 | 618 | 1448 | 1725 |

**Supplementary Table 15.**
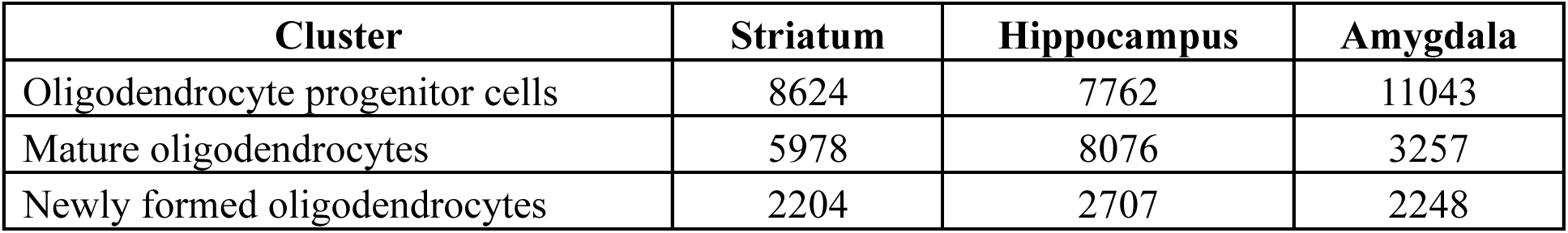
Cell numbers for all oligodendrocyte subclusters by brain region.

**Supplementary Table 16.** Comparison of fractional anisotropy (FA) across 15 white matter tracts between ZIKV-infected and control infants, including sex-based subgroup.

| Fiber tract |  | Female |  | Male |  | Overall |  |
| --- | --- | --- | --- | --- | --- | --- | --- |
|  |  | Control | ZIKA | Control | ZIKA | Control | ZIKA |
|  |  | (N=7) | (N=6) | (N=5) | (N=6) | (N=12) | (N=12) |
| ccg | Mean | 0.361 | 0.363 | 0.352 | 0.364 | 0.357 | 0.364 |
|  | SD | 0.0127 | 0.00571 | 0.0173 | 0.0139 | 0.0147 | 0.0102 |
| cc1d | Mean | 0.368 | 0.375 | 0.358 | 0.382 | 0.364 | 0.378 |
|  | SD | 0.0119 | 0.00889 | 0.0192 | 0.0172 | 0.0154 | 0.0136 |
| cc2d | Mean | 0.44 | 0.428 | 0.414 | 0.44 | 0.429 | 0.435 |
|  | SD | 0.0172 | 0.0164 | 0.0309 | 0.0168 | 0.0263 | 0.0171 |
|  | QC excl | 0 | 1 | 0 | 0 | 0 | 1 |
| cc3d | Mean | 0.378 | 0.371 | 0.344 | 0.36 | 0.364 | 0.366 |
|  | SD | 0.0279 | 0.0184 | 0.0172 | 0.0182 | 0.0288 | 0.0184 |
| ccs | Mean | 0.504 | 0.516 | 0.488 | 0.512 | 0.497 | 0.514 |
|  | SD | 0.0164 | 0.0263 | 0.0402 | 0.0128 | 0.0283 | 0.0198 |
| uf_l | Mean | 0.356 | 0.353 | 0.35 | 0.366 | 0.354 | 0.359 |
|  | SD | 0.0145 | 0.0229 | 0.0174 | 0.00812 | 0.0153 | 0.0177 |
| uf_r | Mean | 0.363 | 0.369 | 0.365 | 0.373 | 0.364 | 0.371 |
|  | SD | 0.0184 | 0.0176 | 0.0182 | 0.00967 | 0.0175 | 0.0137 |
| cg_l | Mean | 0.315 | 0.324 | 0.313 | 0.334 | 0.315 | 0.328 |
|  | SD | 0.0133 | 0.0172 | 0.0223 | 0.00526 | 0.017 | 0.0136 |
|  | QC excl | 1 | 1 | 0 | 2 | 1 | 3 |
| cg_r | Mean | 0.327 | 0.334 | 0.315 | 0.344 | 0.322 | 0.339 |
|  | SD | 0.0142 | 0.0144 | 0.0209 | 0.0116 | 0.0176 | 0.0134 |
|  | QC excl | 0 | 1 | 0 | 1 | 0 | 2 |
| cst_l | Mean | 0.433 | 0.418 | 0.412 | 0.442 | 0.424 | 0.43 |
|  | SD | 0.0162 | 0.0267 | 0.0135 | 0.0257 | 0.0178 | 0.028 |
| cst_r | Mean | 0.434 | 0.41 | 0.406 | 0.441 | 0.422 | 0.425 |
|  | SD | 0.0172 | 0.0191 | 0.0104 | 0.0175 | 0.02 | 0.0237 |
| ilf_l | Mean | 0.331 | 0.338 | 0.321 | 0.336 | 0.327 | 0.337 |
|  | SD | 0.0222 | 0.0262 | 0.0149 | 0.00528 | 0.0195 | 0.018 |
| ilf_r | Mean | 0.347 | 0.355 | 0.339 | 0.358 | 0.344 | 0.356 |
|  | SD | 0.0153 | 0.0178 | 0.0159 | 0.0119 | 0.0153 | 0.0145 |
| fornix_l | Mean | 0.3 | 0.301 | 0.292 | 0.316 | 0.296 | 0.309 |
|  | SD | 0.0269 | 0.0183 | 0.0198 | 0.0112 | 0.0236 | 0.0164 |
| fornix_r | Mean | 0.285 | 0.291 | 0.287 | 0.31 | 0.286 | 0.301 |
|  | SD | 0.0202 | 0.011 | 0.016 | 0.0129 | 0.0178 | 0.015 |

